# Cryo-EM structure of the polyphosphate polymerase VTC: Coupling polymer synthesis to membrane transit

**DOI:** 10.1101/2023.01.27.525886

**Authors:** Wei Liu, Jiening Wang, Véronique Comte-Miserez, Mengyu Zhang, Xuejing Yu, Qingfeng Chen, Andreas Mayer, Shan Wu, Sheng Ye

## Abstract

The eukaryotic polyphosphate (polyP) polymerase VTC complex synthesizes polyP from adenosine triphosphate (ATP) and translocates polyP across the vacuolar membrane to maintain an intracellular phosphate (P_i_) homeostasis. To discover how VTC complex solves this fundamental aspect, we determined a cryo-electron microscopy structure of an endogenous VTC complex (Vtc4/Vtc3/Vtc1) from *Saccharomyces cerevisiae* at 3.1 Å resolution. The structure reveals a heteropentameric architecture of one Vtc4, one Vtc3 and three Vtc1 subunits. The transmembrane region forms a polyP selective channel, probably adopting a resting state conformation, in which a latch-like, horizonal helix of Vtc4 limits the entrance. The catalytic Vtc4 central domain locates on top of the pseudo-symmetric polyP channel, creating a strongly electropositive pathway for nascent polyP that can couple synthesis to translocation. The SPX domain of Vtc4 positively regulates polyP synthesis and regulation of VTC complex. The non-catalytic Vtc3 regulates VTC through a phoshorylatable loop. Our findings, along with the functional data, allow us to propose a mechanism of polyP channel gating and VTC complex activation.

## INTRODUCTION

Phosphate (P_i_) homeostasis is a tightly regulated process in all organisms. Cells face major changes in demand and supply of P_i_, with the strongest one during the S-phase when DNA is duplicated. While cells must meet these P_i_ demands, at the same time, they must safeguard themselves against an excessive cytoplasmic P_i_ concentration. How cells maintain an intracellular P_i_ homeostasis is a fundamental question and of growing interest for medicine and agriculture. Dysfunction of P_i_ homeostasis leads to neurodegeneration of renal Fanconi Syndrome in humans (Ansermet, Moor et al., 2017, Legati, Giovannini et al., 2015), severe growth retardation and dwarfism in plants (Liu, Yang et al., 2015, Puga, Mateos et al., 2014), and lethality in microorganisms (Sethuraman, Rao et al., 2001).

To achieve a delicate balance between the biosynthetic requirements for P_i_ and the risks of an excessive cytoplasmic P_i_ concentration, unicellular organisms maintain important P_i_ stores in membrane-bound, acidocalcisome-like organelles in the form of inorganic polyphosphates, a polymer of up to a thousand P_i_ units linked through phosphoric anhydride bonds (Docampo & Huang, 2016). PolyP influences numerous processes in eukaryotes, ranging from activation of inflammatory responses, wound healing and blood clotting (Gerasimaite & Mayer, 2016, Hassanian, Dinarvand et al., 2015, Hoac, Kiffer-Moreira et al., 2013, Holmstrom, Marina et al., 2013, Mailer, Hanel et al., 2019, Moreno-Sanchez, Hernandez-Ruiz et al., 2012, Schepler, Neufurth et al., 2022, Smith & Morrissey, 2014) to regulation of bone calcification, cation acquisition (Klompmaker, Kohl et al., 2017), protein polyphosphorylation (Azevedo, Livermore et al., 2015, Azevedo, Singh et al., 2018, Bentley-DeSousa, Holinier et al., 2018, Bondy-Chorney, Abramchuk et al., 2020), protein folding (Gray, Wholey et al., 2014), osmoregulation (Lander, Ulrich et al., 2013, Rohloff & Docampo, 2008) and virulence of a series of pathogens (Ikeh, Ahmed et al., 2017). PolyP can also have a major impact on cytosolic Pi homeostasis. Dysregulation of its synthesis can drive cells into Pi starvation or a state of Pi excess (Desfougeres, Gerasimaite et al., 2016). In case of sudden P_i_ starvation, polyP from acidocalcisome-like vacuoles can guarantee sufficient P_i_ reserves to finish the next cell cycle and make an ordered transition into a robust quiescent state. PolyP also buffers transient spikes in P_i_ consumption, which can occur during S-phase (Bru, Martinez-Lainez et al., 2016).

In prokaryotes, the polyphosphate kinase PPK1/2 catalyzes the γ-phosphate transfer of ATP to produce polyP chains (Akiyama, Crooke et al., 1992). Despite the widespread presence of polyP and acidocalcisome-like vacuoles in eukaryotes, only a single eukaryotic polyP-synthesizing enzyme could so far be isolated. This vacuolar transporter chaperone (VTC) complex, originally identified in yeast but with homologs in a wide variety of lower eukaryotes, has provided insights into the mechanisms underlying the polyP synthesis (Hothorn, Neumann et al., 2009). The aim of this study is to address three fundamental questions related to VTC complex. The first question is related to the stoichiometry and the assembly of native VTC complexes. VTC complexes of *Saccharomyces cerevisiae* contain four subunits: Vtc1, Vtc2, Vtc3 and Vtc4 (Cohen, Perzov et al., 1999, Muller, Bayer et al., 2002). Vtc1 is a small membrane protein only containing three transmembrane helices. Vtc2, Vtc3 and Vtc4 are highly homologous in sequence, share a similar transmembrane domain with Vtc1 at the C-terminus, and have an N-terminal SPX (*<u>S</u>*yg1/*<u>P</u>*ho81/*<u>X</u>*PR1) domain that plays key role in P_i_ homeostasis (Wild, Gerasimaite et al., 2016), and a tunnel-shaped, central domain (Hothorn et al., 2009). The central domain of Vtc4 is a polyP polymerase that synthesizes polyphosphate using ATP as a substrate, while that of Vtc2 or Vtc3 is catalytically inactive (Hothorn et al., 2009, Wild et al., 2016). The crystal structures of the central domains of Vtc2 and Vtc4, and the SPX domain of Vtc4 had been determined (Hothorn et al., 2009). However, these structures provide limited information about the stoichiometry and the assembly of VTC complexes. The second question is related to the functional integrity of VTC complexes. The VTC complex is not only a polyP polymerase but also a polyP translocase. To avoid the toxicity of the accumulation of polyP in the cytoplasm, polyP synthesis and the immediate translocation of polyP into the vacuole are coupled (Gerasimaite, Sharma et al., 2014, McCarthy, Abramchuk et al., 2022). However, how they are coupled remains unclear. The third question is related to the regulation of VTC complexes.

When cytosolic P_i_ concentration is sufficiently high, VTC complexes should synthesize polyP efficiently. While with low cytosolic P_i_ concentration, VTC complexes should be switched off to maintain a physiological P_i_ concentration. The activity of VTC complexes is regulated through inositol-based signaling molecules, including the highly phosphorylated, diffusible inositol polyphosphates (InsPs) and inositol pyrophosphates (PP-InsPs) (Wild et al., 2016).

To address these questions, we performed functional assays and cryo-EM structural analysis on endogenous *Saccharomyces cerevisiae* VTC complex. The cryo-EM structure, as well as the detailed functional assay, reveal an unexpected heteropentameric architecture, a coupled polyP polymerase and translocase, a positively regulatory SPX domain, and a phosphorylation dependent regulatory loop, and provide insights into the activation and regulation mechanism of the VTC complex, as well as the polyP channel gating mechanism.

## RESULTS

### Purified VTC complex synthesizes polyP in an ATP- and inositol polyphosphate-dependent manner

The VTC complex of *Saccharomyces cerevisiae* contains four subunits: Vtc1, Vtc2, Vtc3 and Vtc4 (Cohen et al., 1999, Muller et al., 2002). We first performed pull-down assays, and confirmed that no interaction exists between Vtc2 and Vtc3, either directly or indirectly (Figure S1A), indicating that there exist two different VTC complexes, Vtc4/Vtc3/Vtc1 and Vtc4/Vtc2/Vtc1, as also revealed previously (Hothorn et al., 2009). Consistently, knockout of VTC1 or VTC4 alone, or of both VTC2 and VTC3, significantly reduced the cellular PolyP content (Figure 1A), indicating that the catalytic subunit Vtc4 is necessary but not sufficient for polyP synthesis *in vivo*. Individual knockout of VTC2 or VTC3, which disrupted the formation of only one of the two VTC complexes, did not significantly reduce the cellular polyP content (Figure 1A). Interestingly, knockout of VTC2 significantly enhanced the cellular polyP content (Figure 1A). This indicates that Vtc4/Vtc3/Vtc1 and Vtc4/Vtc2/Vtc1 complexes independently synthesize polyP, and suggests a compensatory mechanism to boost the function of one complex when the other one loses function for the maintenance of the intracellular polyP content. Given this redundancy, hereafter we measured cellular polyP content in VTC2 knockout cells to exclude the interference of Vtc4/Vtc2/Vtc1 complex and focus the analysis on substitutions in the Vtc4/Vtc3/Vtc1 complex.

**Figure 1.**
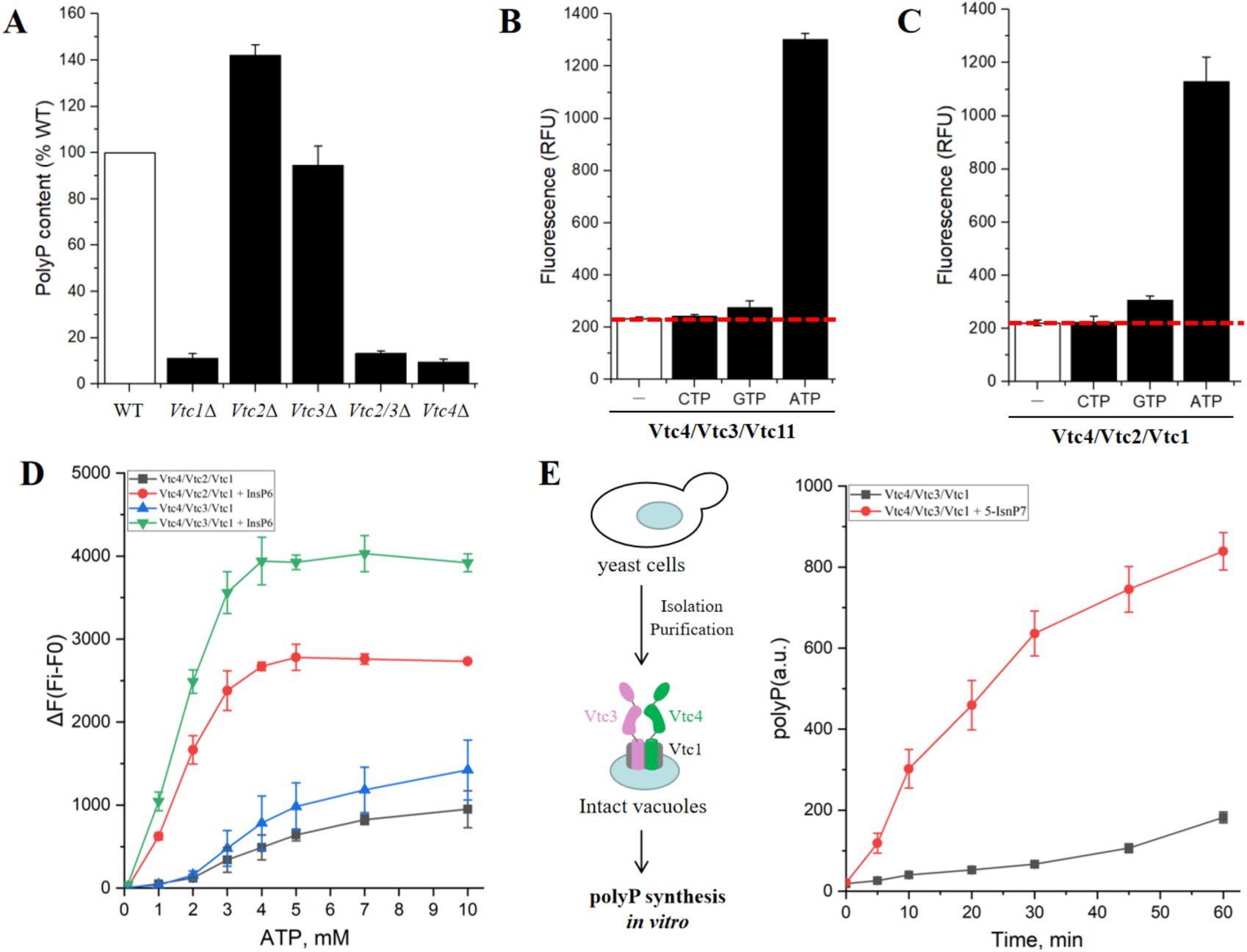
Functional characterization of VTC complexes. (A) PolyP accumulation *in vivo*. The polyP content of wild-type cells was set to 100%. Knockout of Vtc1, Vtc2, Vtc3 or Vtc4 impacts cellular polyP levels. Data show the mean±s.d (n=3). (B and C) Purified endogenous (B) Vtc4/Vtc3/Vtc1 and (C) Vtc4/Vtc2/Vtc1 complexes synthesize polyP from ATP, GTP or CTP *in vitro*. 6 μg of Vtc4/3/1 complex or Vtc4/2/1 complex and 5 mM ATP/GTP/CTP were incubated for 60 min at 4°C, the reaction was stopped by the addition of 15 mM EDTA and 15 μM DAPI, and the fluorescence was measured. Data show the mean±s.d (n=3). (D) PolyP synthesis curves of purified endogenous Vtc4/3/1 and Vtc4/2/1 complexes at different ATP concentrations in the absence or presence of InsP6 *in vitro*. The reaction system is similar to (B) and (C). The concentration of InsP6 was 10 mM. F0 represents the fluorescence value without ATP, as a blank control; Fi represents the fluorescence value at a specific ATP concentration (i=0.1 mM, 1 mM, 2 mM, 3 mM, 4 mM, 5 mM, 7 mM, 10 mM), as the experimental group. ΔF represents the increase in fluorescence value at a specific ATP concentration. Data show the mean±s.d (n=3). (E) PolyP synthesis by isolated vacuoles carrying VTC complexes in the absence or presence of 1μM 5-IP7 *in vitro*. The reaction system is detailed in Methods. Data show the mean±s.d (n=3).

We inserted an affinity tag (His-TEV-Protein A) at the C-terminus of either Vtc3 or Vtc2, and individually purified the endogenous Vtc4/Vtc3/Vtc1 and Vtc4/Vtc2/Vtc1 complexes (Figure S1B, S1C). We performed the polyP synthesis experiments on the intact VTC complexes and observed that both Vtc4/Vtc3/Vtc1 and Vtc4/Vtc2/Vtc1 complexes retain the ability to synthesize polyP from ATP *in vitro* in a divalent cation dependent manner (Figure S2A, S2B), in agreement with a previous study revealing that the central domain (Vtc4^189-480^) of Vtc4 is a polyP polymerase (Hothorn et al., 2009). The synthesized polyP could be degraded by Ppx1, a polyphosphatase in yeast that specifically hydrolyzes polyP (Figure S2A). While ATP, GTP and CTP all interact with the central domain of Vtc4 with binding affinities at micromolar range (Hothorn et al., 2009), polyP synthesis was significantly reduced when ATP was replaced with GTP, an ATP analog sharing a similar purine moiety, and it was completely eliminated upon the replacement of ATP with CTP (Figure 1B, 1C). These data demonstrate that polyP synthesis by the VTC complex is preferentially driven by ATP.

We next measured polyP synthesis as a function of ATP concentration (Figure 1D). VTC activity showed a steep, linear dependence on ATP concentration, reaching saturation at 3-4 mM. With a Km value of around 2 mM ATP, VTC shows a low affinity for ATP, which could be relevant to downregulate VTC activity. This is crucial to the physiological operation of VTC complexes in the cellular context, as free cellular ATP levels in yeast have been estimated to be ∼ 1–2 mM (Ingram & Barnes, 2000, Ozalp, Nielsen et al., 2010, Pluskal, Hayashi et al., 2011). In a situation where P_i_ is abundant and VTC is maximally activated through inositol pyrophosphates (PP-InsPs), the high Km value provides an inbuilt mechanism to downregulate polyP synthesis if ATP supply of the cell runs low. In this way, VTC could integrate intracellular phosphate reception and signaling pathway with the cellular energy status.

In response to cellular P_i_ availability, PP-InsPs increase a polyP accumulation. In line with this, InsPs and PP-InsPs can activate VTC (Gerasimaite, Pavlovic et al., 2017, Lonetti, Szijgyarto et al., 2011, Wild et al., 2016). PolyP synthesis by purified VTC 4/3/1 complex was strongly stimulated by InsP6, especially at low concentrations of ATP (Figure 1D). Without the addition of ATP, InsP6 alone did not produce a fluorescence signal (Figure S2C), demonstrating the regulatory role of InsP6.

Given that the PP-InsPs are physiological activators of the VTC complex, and the VTC complex integrated in the intact membrane displays much higher activity than the purified complex (Gerasimaite et al., 2014), we also performed in vitro polyP synthesis experiments using purified vacuoles. As shown in Figure 1E, the VTC complex from purified vacuoles synthesizes polyP in an ATP-dependent manner, and PP-InsPs significantly enhance polyP synthesis by more than 10-fold.

### Overall architecture of yeast Vtc4/Vtc3/Vtc1 complex

To understand the polyP synthesis and transport mechanism, we analyzed the purified Vtc4/Vtc3/Vtc1 complex by single-particle cryo-electron microscopy (Cryo-EM), yielding a structure at an overall resolution of 3.1 Å (Figure S3, S4). The cryo-EM density map was of sufficient quality to allow modeling of almost the entire complex, including the N-terminal SPX domains, the central domains, the C-terminal transmembrane (TM) domains of Vtc4 and Vtc3, and the TM domains of Vtc1 (Figure S5).

The structure of the Vtc4/Vtc3/Vtc1 complex reveals an unexpected heteropentameric architecture with a subunit stoichiometry of one Vtc4, one Vtc3 and three Vtc1 subunits (Figure 2A). The transmembrane domains of Vtc1, Vtc3 and Vtc4, which share approximately 15% amino acid sequence identity, adopt similar backbone conformations (Figure 2B). Three transmembrane helices (TM1-TM3) from each subunit assemble in a pseudo-symmetrical fashion forming a cylinder-shaped pentameric transmembrane domain. When viewed from the cytoplasmic side, the arrangement of subunits around the transmembrane domain is Vtc3-Vtc4-Vtc1-Vtc1- Vtc1 in a counter-clockwise direction (Figure 2C). The arrangement results in a different subunit packing environment for each Vtc1. We therefore refer to the three Vtc1 subunits as Vtc1(α), Vtc1(β) and Vtc1(γ) for clarity (Figure 2C). The TM1 helices from each subunit form an inner ring lining a pore that tapers as it traverses towards the intravacuolar side of the membrane. The TM2 and TM3 helices from each subunit form an outer ring surrounding the inner ring, which forms a central pore for polyP translocation into the vacuole. Compared to Vtc1 or Vtc4, Vtc3 has an additional, amphipathic MX helix at the C-terminus (Figure S6A). The MX helix of Vtc3 runs parallel to the vacuolar face of the membrane, forming hydrophobic interactions with TM2 and TM3 helices of Vtc4, and TM3 helix of Vtc3 (Figure S6B). Given that Vtc2 and Vtc3 share a same MX helix with high sequence identity (Figure S7), the MX helix of Vtc2 likely adopts the same conformation and performs similar function in the Vtc4/Vtc2/Vtc1 complex in comparison with that of Vtc3 in the Vtc4/Vtc3/Vtc1 complex.

**Figure 2.**
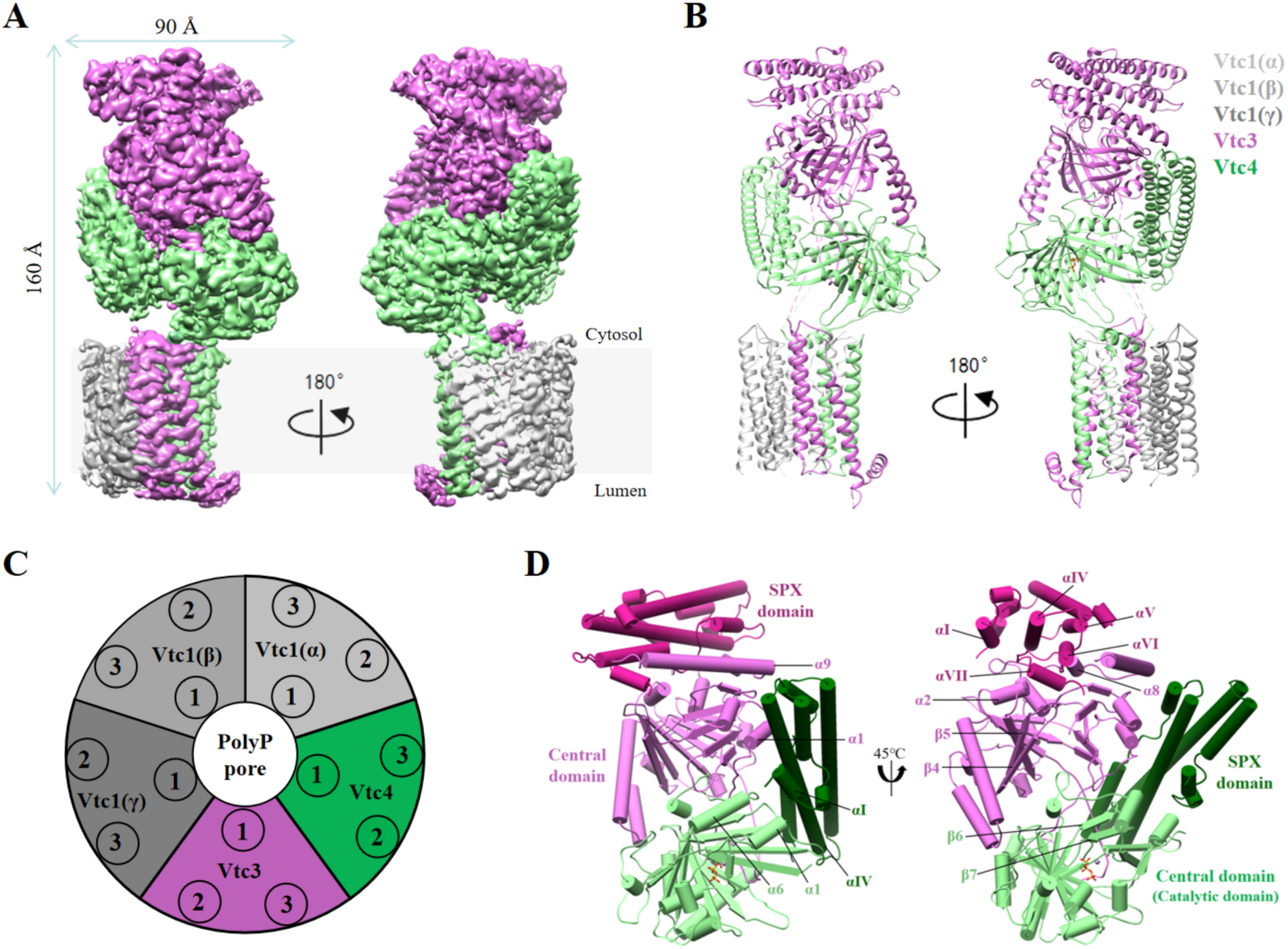
Structure of the yeast Vtc4/Vtc3/Vtc1 complex. (A) Cryo-EM 3D map of the Vtc4/Vtc3/Vtc1 complex, showing front and back views. Color codes for the subunits of the complex are indicated. (B) An atomic model shown in cartoon and colored as in A. The triphosphate and Mn^2+^ are shown in orange and brown, respectively. (C) Top view of the model of Vtc4/Vtc3/Vtc1 complex. The numbers 1, 2 and 3 represent TM1, TM2 and TM3, respectively, where TM1 is at the N-terminus of the sequence and TM3 is at the C-terminus of the sequence. (D) Structure of the asymmetrically arrangement of the intracellular region of the Vtc4/Vtc3/Vtc1 complex. In order to distinguish, the SPX domain and central domain of Vtc3 are shown in violet red and orchid, respectively; the SPX domain and central domain of Vtc4 are shown in dark green and light green, respectively.

The structure of the Vtc4/Vtc3/Vtc1 complex reveals an asymmetrical arrangement of the cytosolic region in contrast to the relatively symmetrical arrangement of the transmembrane region. Vtc4, Vtc3 and Vtc2 all contain an N-terminal SPX domain, a central domain, and a C-terminal transmembrane domain (Hothorn et al., 2009, Wild et al., 2016). The central domain (Vtc4^189-480^) of Vtc4 is a polyP polymerase, while those of Vtc2 and Vtc3 are catalytically inactive, likely playing an accessory function (Hothorn et al., 2009). Correspondingly, the catalytically active central domain of Vtc4 is the only cytosolic domain that interacts with the transmembrane pore (Figure S5). The catalytically inactive central domain of Vtc3 stacks on top of the central domain of Vtc4 in a head to tail manner, forming a heterodimer (Figure 2D). Interestingly, the two SPX domains adopt different positions relative to the respective central domains. The SPX domain of Vtc4 locates at one side of the central domain of Vtc4, using its α1 and α3 helices to interact with the α1 and α6 helices and the β6 and β7 strands of the central domain of Vtc4 (Figure 2D).

By contrast, the SPX domain of Vtc3 locates at the other side of the central domain of Vtc3, using a different set of α helices (α1, α4, α5, α6 and α7) to interact with three α helices (α2, α8 and α9) and two β strands (β4 and β5) of the central domain of Vtc3 (Figure 2D). The asymmetrical arrangement of the cytosolic region of the Vtc4/Vtc3/Vtc1 places both SPX domains close to each other on the same side of the complex (Figure 2D). The SPX domain of Vtc4 not only interacts with the central domain of Vtc4, but also interacts with two α helices (α1 and α9) of the central domain of Vtc3 (Figure 2D). By contrast, the SPX domain of Vtc3 only interacts with the central domain of Vtc3 (Figure 2D).

### PolyP channel in a resting and fastened state

The transmembrane domain of the Vtc4/Vtc3/Vtc1 complex resembles a cylinder formed from five subunits in a pseudo-symmetrical arrangement about a central axis. The pore is lined by five TM1 helices, forming ‘rings’ of positively charged residues. The cytoplasmic vestibule of the VTC channel contains two positively charged rings, with K24 of Vtc1, K698 of Vtc3, and K622 of Vtc4 forming one, and R31 of Vtc1, R705 of Vtc3, and R629 of Vtc4 forming the other one, rendering the surface strongly electropositive (Figure 3A, 3B). These positively charged rings may constitute a polyP selectivity filter in the vestibule of the VTC channel, and probably serve to attract polyP to the channel mouth. To test the role of these rings for polyP synthesis and translocation, we created 6 charge-reversing point mutations. All these mutations significantly reduced cellular polyP content (Figure S8A).

**Figure 3.**
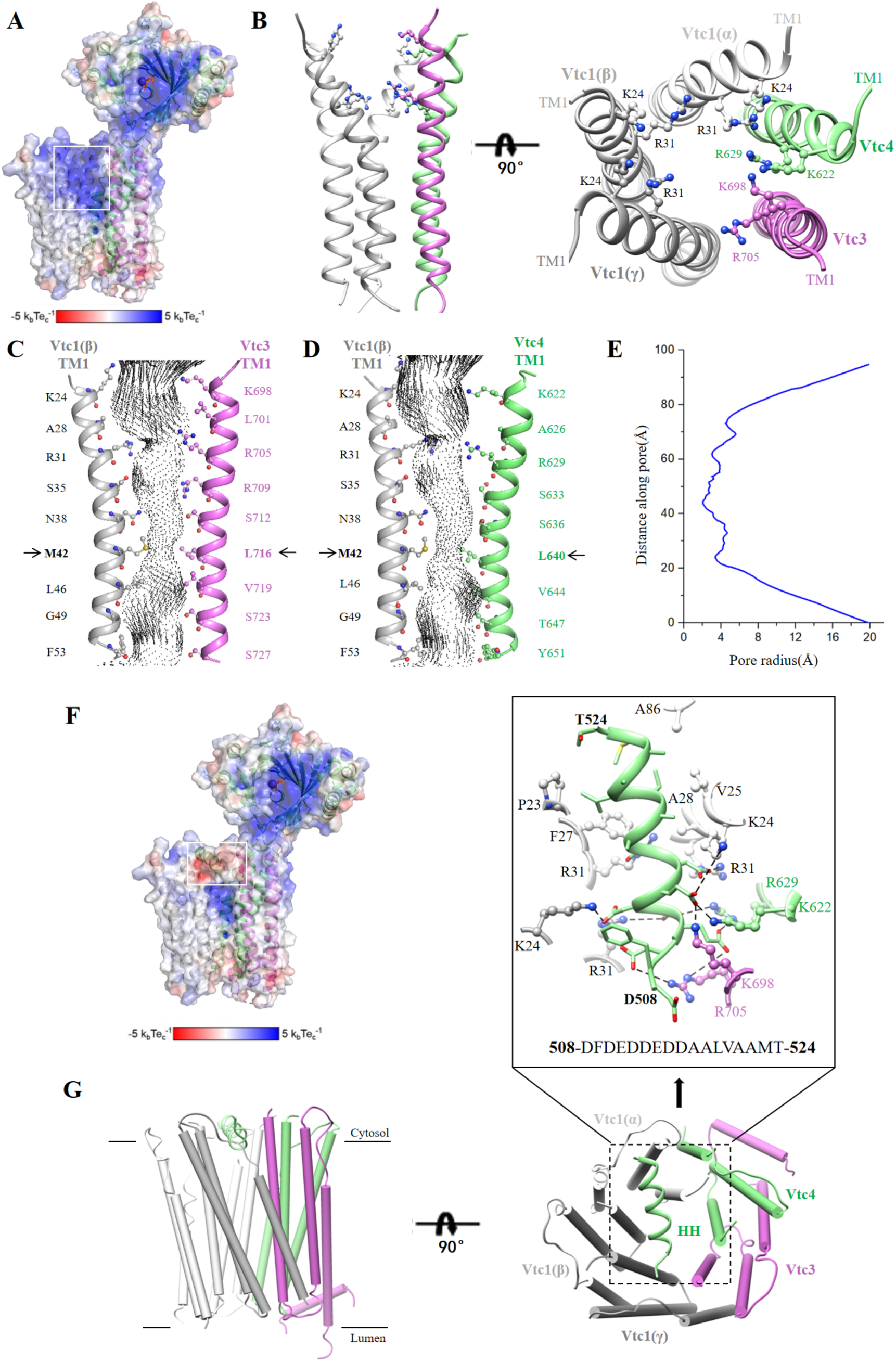
Conductance and permeation pore structure of the Vtc4/Vtc3/Vtc1 complex. (A) Cutaway of the Vtc4/3/1 complex showing electrostatic surface potential along the polyP-conducting pathway, excluding the horizontal helix, HH. The transparency of the electrostatic surface potential is set to 0.5. (B) Side and top view of the structure of TM1 of Vtc4/Vtc3/Vtc1 complex. The cytoplasmic vestibule of the VTC channel contains two positively charged rings, with K24 of Vtc1, K698 of Vtc3, and K622 of Vtc4 forming one, and R31 of Vtc1, R705 of Vtc3, and R629 of Vtc4 forming the other one。 (C) TM1 α-helices from opposing Vtc1(β) and Vtc3 subunits with side chains shown for pore-lining residues. Spheres represent the solvent-accessible volume of the polyP channel. The black arrow points to the narrowest point of the channel. (D) TM1 α-helices from opposing Vtc1(β) and Vtc4 subunits with side chains shown for pore-lining residues. Spheres represent the solvent-accessible volume of the polyP channel. The black arrow points to the narrowest point of the channel. (E)Profile of pore radius of the Vtc4/Vtc3/Vtc1 complex. (F)Cutaway of the Vtc4/3/1 complex showing electrostatic surface potential along the polyP-conducting pathway, including the horizontal helix, HH. The transparency of the electrostatic surface potential is set to 0.5. (G) Side and top view of the structure of transmembrane helices of Vtc4/Vtc3/Vtc1 complex. Horizontal helix, HH (508-DFDEDDEDDAALVAAMT-524).

The TM1 helices taper inwards from the cytosolic side to the intravacuolar side, with a ring of hydrophobic residues, including M42 of Vtc1, L716 of Vtc3, and L640 of Vtc4, defining the narrowest point, just 4 Å in diameter (Figure 3C, 3D, 3E). Since this point is too narrow to permit the passage of polyP (with a Pauling radius of 3 Å) the structure likely represents a non-conducting resting state of the pore.

In addition, the entrance of the PolyP channel is fastened by a latch-like, horizonal helix (HH) (Figure 3F, 3G). This helix is part of a linker of over hundred residues connecting the central domain and the transmembrane domain of Vtc4 (Figure S7). The majority of the residues are hydrophilic, without density in the cryo-EM map, indicating that the linker is highly flexible. However, the N-terminal part of the linker (^508^DFDEDDEDDAALVAAMT^524^), which is rich in acidic residues, forms an α helix horizontally latching the entrance of the transmembrane channel (Figure 3G). The N-terminus of the helix nestles at the Vtc1(γ)-Vtc3 interface, with the acidic residues forming multiple salt bridges with the positively charged residues at the channel mouth (Figure 3G). The C-terminus of the helix forms multiple hydrophobic interactions with Vtc1(α) and Vtc1(β) (Figure 3G). To probe the importance of this horizonal helix, we deleting it (residues 508-524) from Vtc4 and observed an approximately 20% higher cellular polyP content in the respective mutant (Figure S8B).

### Coupled polyP synthesis and translocation

The structure of the Vtc4/Vtc3/Vtc1 complex clearly supports the concept of a coupled polyP polymerase and translocase. The structures of the two central domains of the Vtc4/Vtc3/Vtc1 complex are highly similar, with a r.m.s. deviation of 1.8 Å for 276 Cα atoms. Both central domains contain a central tunnel formed by antiparallel β strands, with the majority of the α helices franking the tunnel wall. The α helix (α7) at the C-terminus of the central domain of Vtc4 forms a “helical plug” at one end of the tunnel, reducing the tunnel to a small size only allowing polyP to pass (Figure S9).

The central domain of Vtc3 contains two additional α helices (α8 and α9) at the C-terminus (Figure S9). These additional α helices, together with the SPX domain of Vtc3, completely block one end of the tunnel of the central domain of Vtc3, likely rendering it inactive (Figure S9). The tunnel walls are lined by conserved basic residues (Figure 4A). Confirming it role as the catalytically active subunit of a polyP polymerase, the Vtc4 central domain shows an endogenous Mn^2+^-bound triphosphate in the tunnel center (Figure S10). The Mn^2+^ is chelated by carboxylate oxygens of E426 and the triphosphate oxygens in a distorted square-based pyramidal configuration (Figure 4A). The triphosphate is coordinated by six conserved basic residues (K200, R264, R266, K281, R361, K458), a serine (S457) and a tyrosine (Y359) (Figure 4A). These residues are critical for polyP synthesis, as alanine mutations of K200, R264, R266, K281, R361, K458, S457, and the phenylalanine mutation of Y359, all significantly reduce polyP content of respective mutant cells (Figure 4B). Structure based sequence alignment revealed that among these residues, the only difference is K458 of Vtc4, which is I522 in Vtc2, and L527 in Vtc3 (Figure S11A, S11B, S11C, S11D). Substituting K458 of Vtc4 to either leucine or isoleucine significantly reduced polyP content, underlining the critical role of K458 for polyP synthesis (Figure 4B).

**Figure 4.**
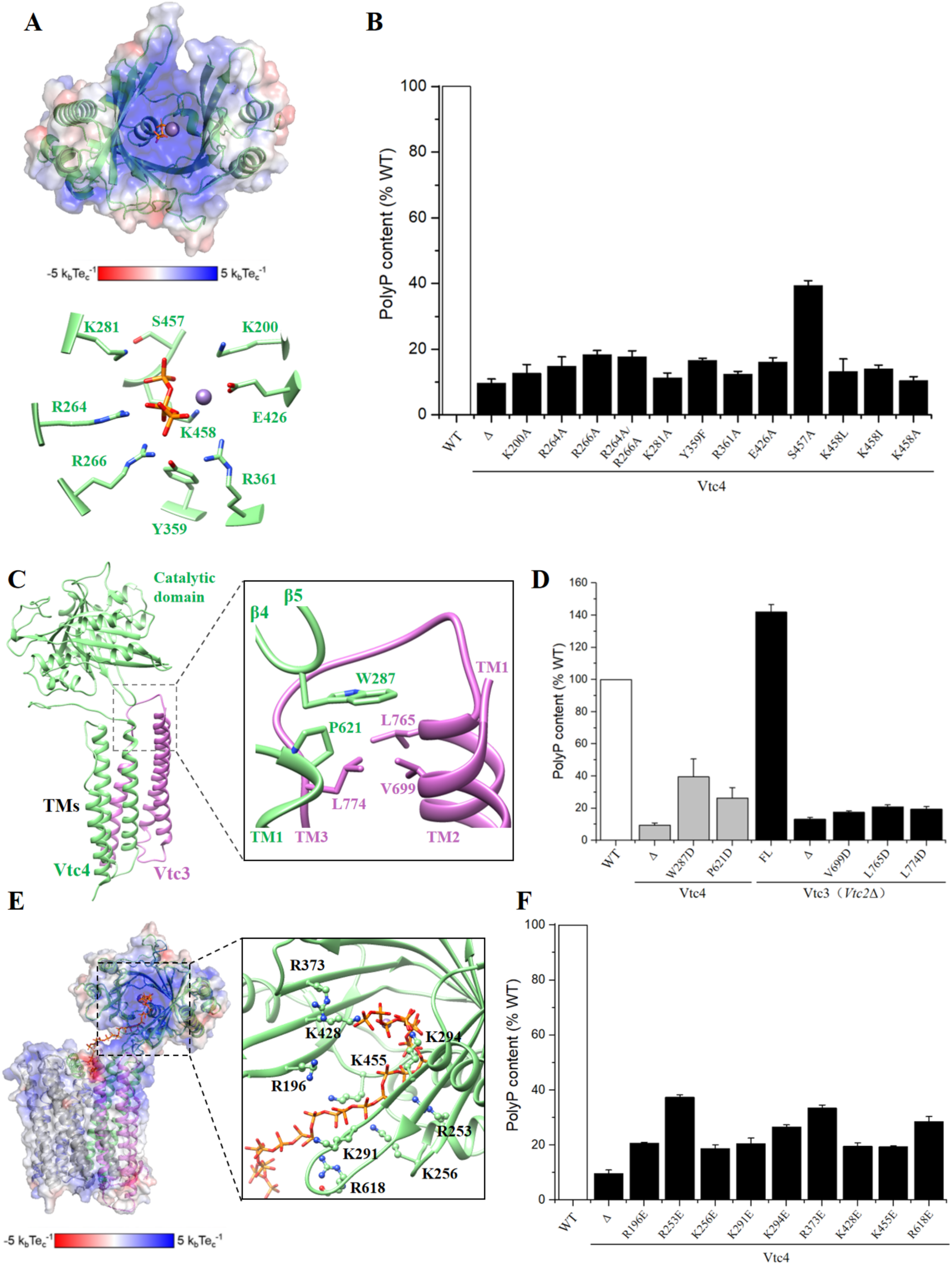
Structural and functional data of the VTC complex reveal that polyP synthesis and transport are coupled. (A) Structure and electrostatic surface potential of the central domain of Vtc4. The triphosphate and Mn^2+^ are shown in orange and brown, respectively. Some key residues are shown. (B) Cellular polyP content of Vtc4p point mutants expressed under the control of their native promoters in the *vtc4Δ* backgrounds. Data show the mean±s.d (n=3). (C) Interactions between the β4-β5 loop of Vtc4 and the transmembrane domains of Vtc3 and Vtc4. (D) Cellular polyP content of Vtc4p and Vtc3p point mutants expressed under the control of their native promoters in the *vtc4Δ* and *vtc3Δ*(*vtc2Δ*) backgrounds, respectively. Data show the mean±s.d (n=3). (E) Superposition of the central domain of Vtc4 and the central domain of the polyP- bound Vtc4(PDB: 3G3Q) structures. The structure of the central domain of polyP- bound Vtc4 is shown in blue. The polyP chains are shown in orange to overlap the triphosphates. (F) Cellular polyP content of Vtc4p point mutants expressed under the control of their native promoters in the *vtc4Δ* backgrounds. Data show the mean±s.d (n=3).

Although the central domain and the transmembrane domain of Vtc4 are covalently connected, the majority of the linker in between is flexible without observable density in the cryo-EM map. The central domain of Vtc4 interacts with the transmembrane pore via contacts between the β4-β5 loop of Vtc4 and the TM2-TM3 loop of Vtc3, as well as contacts between the loop before TM1 of Vtc4 and the β4-β5 loop, α2, and the loop after β11 of Vtc4 (Figure 4C). The β4-β5 loop of Vtc4 is extended and protrudes into a hydrophobic pocket formed between the transmembrane domains of Vtc3 and Vtc4, with the aromatic side chain of Trp287 interacting with Val699, Leu774 and Leu765 of the TM2-TM3 loop of Vtc3, and Pro621 of the loop before TM1 of Vtc4 (Figure 4C). To confirm the importance of the observed interactions between the central domain of Vtc4 and the transmembrane pore, we created point mutants designed to disrupt the hydrophobic contact by changing hydrophobic residues to acidic residues, and observed that they all significantly reduce cellular polyP content (Figure 4D).

The interactions bring the catalytically active central domain of Vtc4 in close proximity to the transmembrane pore, with the tunnel walls directly connecting to the vestibule of the pore. Superimposition of the previously determined central domain of Vtc4 (Hothorn et al., 2009) with the one determined in this study reveals that the phosphate polymer overlaps with the triphosphate and winds through the tunnel towards the vestibule of the pore (Figure 4E), suggesting that the nascent polyP travels from the active site of the central domain of Vtc4 to the vestibule of the transmembrane pore, thus translocating across the membrane. In addition, the traveling pathway of polyP is strongly electropositive, which probably feeds the polyP product through the membrane pore into the lumen of the vacuole. To confirm the importance of the electropositive pathway, we created point mutants designed to switch the electrostatic potential by changing positively charged residues to acidic residues and observed that the intracellular content of polyP was significantly reduced (Figure 4F).

### The SPX domain of Vtc4 is critical for polyP synthesis and PP-InsPs regulation

Both Vtc3 and Vtc4 contain an N-terminal SPX domain that may sense the cellular P_i_ levels. The structures of the two SPX domains are highly similar, with a r.m.s. deviation of 1.7 Å for 135 Cα atoms. Both SPX domains share an N-terminal helical hairpin formed by two small helices, αI and αII, and a three-helix bundle formed by two long helices, αIII and αIV, together with two smaller C-terminal helices, αV and αVI (Figure S7). The SPX domain of Vtc3 contains an additional helix, αVII, at the C-terminus (Figure 5A), forming hydrophobic interactions with αIV and αVI helices (Figure 5A). Both SPX domains harbor a positively charged surface formed by multiple conserved lysine residues from helices αII and αIV (Figure S12B, S12C), and can interact with a phosphate-containing ligand with little specificity and selectivity (Wild et al., 2016).

**Figure 5.**
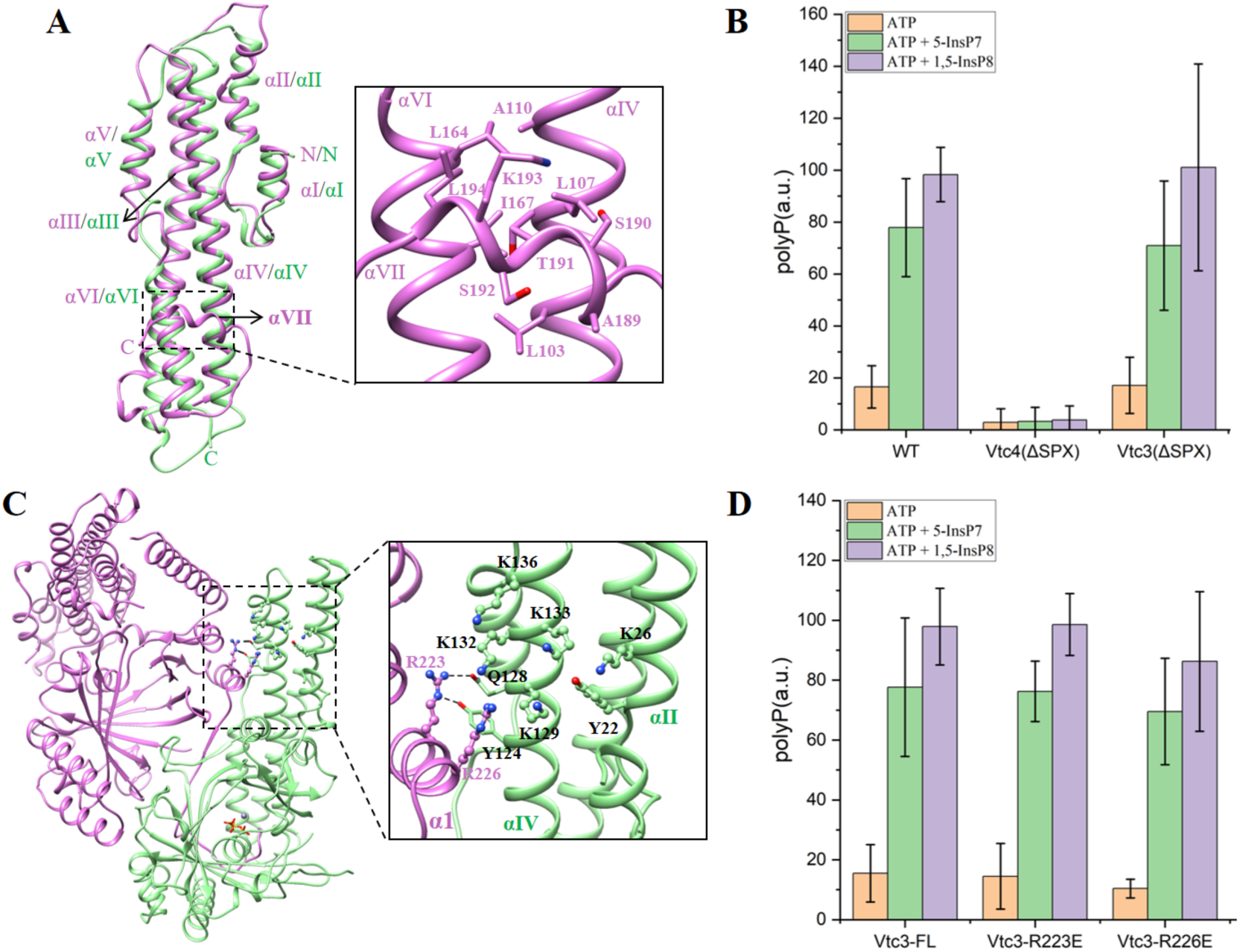
The SPX domain of Vtc4 regulates polyP synthesis in an PP-InsPs- dependent manner. (A) Superposition of the SPX domain of Vtc4 and the SPX domain of Vtc3. The SPX domain of Vtc3 is shown in orchid, and the SPX domain of Vtc4 is shown in light green. (B) PolyP synthesis by isolated vacuoles carrying Vtc4(ΔSPX)/Vtc3/Vtc1 complex, Vtc4/Vtc3(ΔSPX)/Vtc1 complex or Vtc4/Vtc3/Vtc1 complex in the absence or presence of 1 μM 5-IP7 or 1,5-IP8 *in vitro*. The reaction system is detailed in Methods. Data show the mean±s.d (n=3). (C) A conserved arginine residue on Vtc3 is involved in the regulation of InsP6. (D) PolyP synthesis by isolated vacuoles carrying Vtc4/Vtc3(R223E)/Vtc1 complex, Vtc4/Vtc3(R226E)/Vtc1 complex or Vtc4/Vtc3/Vtc1 complex in the absence or presence of 1 μM 5-IP7 or 1,5-IP8 *in vitro*. The reaction system is detailed in Methods. Data show the mean±s.d (n=3).

An interesting aspect of the Vtc4/Vtc3/Vtc1 complex is that the SPX domain of Vtc4 interacts with both the central domain of Vtc3 and Vtc4 while the SPX domain of Vtc3 only interacts with the central domain of Vtc3. To probe the role of the two SPX domains, we individually truncated the SPX domain of Vtc3 or Vtc4 and performed polyP synthesis experiments on the purified mutant Vtc4/Vtc3/Vtc1 complexes (Figure S13A, S13B). Truncation of the SPX domain of Vtc3 reduced polyP synthesis activity of the complex only slightly, and preserved the stimulation of its activity by InsP6 (Figure S13C), indicating that the SPX domain of Vtc3 is not essential for stimulation of polyP synthesis by InsP6. By contrast, truncation of the SPX domain of Vtc4 significantly impaired polyP synthesis activity of the Vtc4/Vtc3/Vtc1 complex (Figure S13C) and addition of InsP6 reduced this activity further instead of stimulating it (Figure S13C). Thus, the SPX domain of Vtc4 is critical for polyP synthesis and InsP6 regulation. The isolated vacuoles of the mutant Vtc4/Vtc3/Vtc1 complex with the truncation of the SPX domain of Vtc4 completely lose the polyP synthesis activity. And the addition of 5-IP7 or 1,5-IP8 no longer restores the polyP synthesis activity (Figure 5B). While the isolated vacuoles of the mutant Vtc4/Vtc3/Vtc1 complex with the truncation of the SPX domain of Vtc3 generate less polyP than that of wild type, and the addition of IP7 or 1,5-IP8 enhances the polyP synthesis activity (Figure 5B). Taken together, the data of purified complex and isolated vacuole all demonstrate that the SPX domain of Vtc4 is critical for polyP synthesis and PP-InsPs regulation.

The structure of the Vtc4/Vtc3/Vtc1 complex reveals that the positively charged surface of the SPX domain of Vtc4 is close to the α1 helix of Vtc3 (Figure 5C). Two arginine residues of the α1 helix of Vtc3, R223 and R226, are strictly conserved between Vtc2 and Vtc3 (Figure S7). To probe the potential InsP6 regulation role of the conserved arginine residues, we created point mutants by changing the basic residues to acidic residues and performed the polyP synthesis experiments on the purified mutant Vtc4/Vtc3/Vtc1 complexes (Figure S14A, S14B). Compared with the wild type complex, both mutant complexes display significantly reduced polyP synthesis in the absence of stimulation by IP6, with a 50% reduced activity for the R226E mutant (Figure S14C). However, addition of InsP6 strongly stimulated also the mutant complexes, conferring them only 10-20% lower activity that the wildtype complexcould not restore the same polyP synthesis activities as the wild type (Figure 5D), suggesting that both residues are involved in the polyP synthesis activity and the InsP6 regulation. The isolated vacuoles of the mutant Vtc4/Vtc3/Vtc1 complex also showed similar results to purified complexes, with the mutant R226E showing stronger effects than that of the mutant R223E (Figure 5D). The addition of IP7 or 1,5-IP8 enhances the polyP synthesis activity (Figure 5D), further suggesting that both residues are also involved in the PP-InsPs regulation.

### A regulatory loop of VTC3

An interesting aspect of the Vtc4/Vtc3/Vtc1 complex is the conformation and orientation of a loop between α1 and β2 of the central domain of Vtc3. In comparison with Vtc4, this loop of Vtc3 is unusually long, containing over sixty amino acids (Figure S7). The N-terminal half of the loop (^228^LPALVYASVPNENDDFVDNLES D^250^), which is rich in acidic residues, forms of a nine-residue loop (^228^LPALVYASV^236^), a four-residue turn (^237^PNEN^240^), a five-residue loop (^241^DDFVD^245^) and a five-residue turn (^246^NLESD^250^). It winds across the heterodimeric interface between the two central domains and the tunnel exit of the Vtc4 central domain, interacting with β1, α1, β2, β5, α4, α5 and β7 of Vtc4 (Figure 6A). The last five-residue turn loop (^246^NLESD^250^) of the loop is close to the triphosphate observed in the structure and forms multiple interactions with the positively charged residues of Vtc4, including R196, R280, K281, K294, K300, R373 and K428, suggesting a regulatory role on polyP synthesis of this loop (Figure 6A).

**Figure 6.**
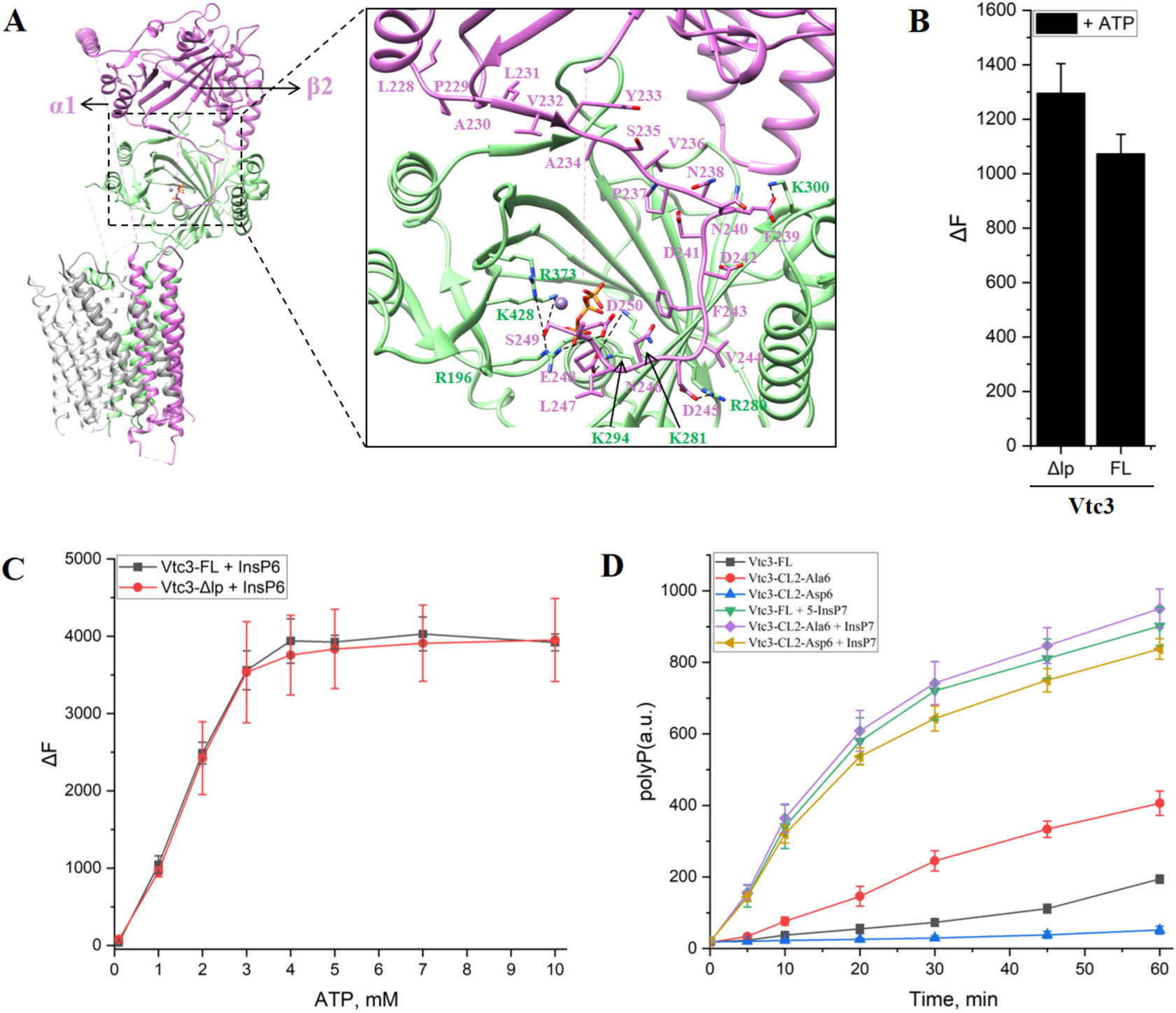
Structure and function of the regulatory loop of Vtc3. (A) Structure of the regulatory loop of Vtc3. The loop is located between α1 and β2 of Vtc3, and the loop sequence consists of ^228^LPALVYASVPNENDDFVDNLESD^250^. Interactions (dotted lines) are shown. (B) Purified truncated Vtc4/Vtc3(Δlp)/Vtc1 complexes synthesize polyP in the presence ATP *in vitro*. Δlp indicates that the regulatory loop of Vtc3 (residues 234- 292) was replaced by a small linker (GGSGGS). The reaction system is detailed in Methods. Data show the mean±s.d (n=3). (C) PolyP synthesis curves of purified endogenous Vtc4/3/1 and truncated Vtc4/Vtc3(Δlp)/Vtc1 complexes at different ATP concentrations in the absence or presence of InsP6 *in vitro*. The reaction system is detailed in Methods. Data show the mean±s.d (n=3). (D) PolyP synthesis by isolated vacuoles carrying Vtc4/Vtc3(CL2-Ala6)/Vtc1 complex, Vtc4/Vtc3(CL2-Asp6)/Vtc1 complex or Vtc4/Vtc3/Vtc1 complex in the absence or presence of 1 μM 5-IP7 or 1,5-IP8 *in vitro*. The reaction system is detailed in Methods. Data show the mean±s.d (n=3).

In an effort to probe the regulatory role of the loop, we first generated a truncated form of Vtc3 by replacing this long loop (residues 234-292) with a short linker (GGSGGS), and performed polyP synthesis experiments on the purified mutant Vtc4/Vtc3/Vtc1 complex (Figure S15A, S15B). However, the mutant Vtc4/Vtc3/Vtc1 complex retained a slightly higher polyP synthesis activity than that of the wild type complex, suggesting a potentially negative regulatory role of the loop (Figure 6B). In addition, the addition of the InsP6 also significantly enhanced polyP synthesis on the mutant Vtc4/Vtc3/Vtc1 complex even at low concentrations of ATP, similar to that of wild type complex (Figure 6C). The moderate effect of the mutant Vtc4/Vtc3/Vtc1 complex led us to probe another possibility, in which we noticed that the C-terminal half of the loop without visible EM density (^251^VRVQPEARLNIG**S**K**S**N**S**L**SS** DGN**S**NQDVEIGK**S**KSVIFPQ**S**Y^292^) contains a cluster of phosphorylation sites, suggesting possible regulation by phosphorylation. To mimic a non-phosphorylated or a phosphorylated state of the loop, we substituted six of its serine residues (S263, S265, S267, S269, S270, S274) by either alanine or aspartate and performed the polyP synthesis experiments with vacuoles purified from respective mutants. In the absence of stimulating PP-InsP, vacuoles carrying the alanine-substituted loop displayed more than 200% higher polyP synthesis activity *in vitro* than vacuoles from wild type, whereas the phospho-mimetic aspartate substituted form had a significantly reduced activity (Figure 6D). Addition of PP-InsP significantly enhanced polyP synthesis of both non-phosphorylated and phosphomimetic forms, conveying similar activity as for the wild type form (Figure 6D). This confirms the negative regulatory role of the loop, silencing the complex when P_i_ (and hence PP-InsPs) are low. This silencing role may be enhanced by its phoshorylation. When P_i_ becomes abundant, this negative regulatory may be overridden by the increase in PP-InsPs. Then, this loop might serve to enhance the dynamic range over which VTC can be regulated, supporting a complete shut-off of polyP synthesis under Pi starvation. Given that Vtc2 and Vtc3 share a same loop with high sequence identity (Figure S7), the loop of Vtc2 likely adopts the same conformation and orientation and performs similar regulatory function as the loop in Vtc3.

### Mechanics of polyP channel gating

Exactly how the TM1 helices move to open the polyP channel of the VTC complex is a difficult question to answer in the absence of direct observation of the open polyP channel structure. However, the polyP channel in such a resting and fastened state provides valuable insight into the conformational changes that apparently underlie polyP channel gating. First, the five-fold symmetry of the polyP channel is broken, as revealed by the two distorted pentagons by connecting the adjacent Cαs of the two positively charged rings at cytoplasmic vestibule of the VTC channel (Figure 7A). The inter-subunit interfaces are extensive, and involve different subunits of the VTC channel. We define principal (+) and complementary (-) subunits and interfaces (Figure 7B), where the principal (+) interface is made up of TM1 and TM3 of the principal subunit, while the complementary (-) interface is made up of TM1 and TM2 of the complementary subunit (Figure 4B). We superimposed the principal subunit of all the five inter-subunit interfaces of the polyP channel, Vtc1(α)- Vtc1(β), Vtc1(β)-Vtc1(γ), Vtc1(γ)-Vtc3, Vtc3-Vtc4 and Vtc4-Vtc1(α), and observed that the two transmembrane domains forming the interface can have various relative positions, with Vtc1(β)-Vtc1(γ) and Vtc3-Vtc4 packing tightly, Vtc1(γ)-Vtc3 loosely, and Vtc1(α)-Vtc1(β) and Vtc4-Vtc1(α) in between (Figure 7B). Correspondingly, the Vtc3-Vtc4 interface buries the most (4150 Å^2^) protein surface area from solvent, followed by Vtc1(β)-Vtc1(γ) (2920 Å^2^), Vtc4-Vtc1(α) (2870 Å^2^) and Vtc1(α)- Vtc1(β) (2660 Å^2^) interfaces. The Vtc1(γ)-Vtc3 interface has the smallest amount of buried surface area (2350 Å^2^). This agrees with the observed asymmetry of the VTC channel. Several reasons can account for the altered relative positioning at the inter-subunit interfaces: forces imposed by the latch-like, horizonal helix of Vtc4, and the constraint of interacting with the central domain of Vtc4. The important point is, however, that the inter-subunit interface is clearly flexible.

**Figure 7.**
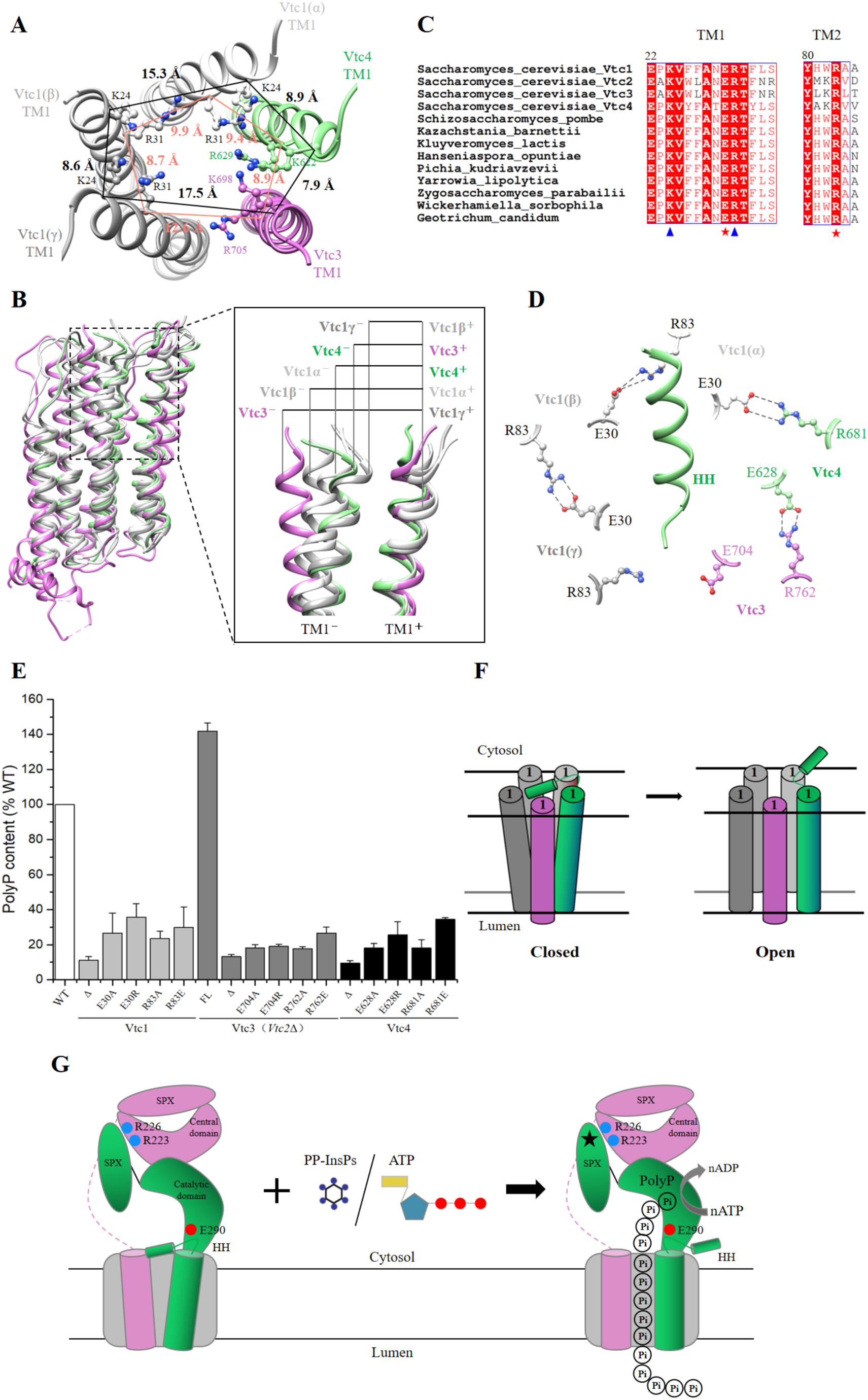
Asymmetric polyP channel and inter-subunit ionic locks. (A) Asymmetry in the channel at the level of the activation and desensitization gates. Residues at the polyP selectivity filter are shown in ball-and-stick representation. Distance between Cα of polyP selectivity filter residues are given in Å. (B) Superposition of the principal subunit of all the five inter-subunit interfaces of polyP channel。 (C) Sequence alignment of Vtc1, Vtc2, Vtc3 and Vtc4 from different species. Protein sequence number in NCBI: *Saccharomyces_cerevisiae*_Vtc1 (ID: NP_010995.1); *Saccharomyces_cerevisiae*_Vtc2 (ID: KZV11596.1); *Saccharomyces_cerevisiae*_Vtc3 (ID:KZV07497.1); *Saccharomyces_cerevisiae*_Vtc4 (ID:QHB096 08.1); *Schizosaccharomyces_pombe* (ID:NP_595683.1); *Kazachstania_barnettii* (ID:XP_041404278.1); *Kluyveromyces_lactis* (ID: QEU59996.1); *Hanseniaspora_opuntiae* (ID: OEJ89736.1); *Pichia_kudriavzevii* (ID:ONH77772.1); *Yarrowia_lipolytica* (ID: QNP96953.1); *Zygosaccharomyces_parabailii* (ID:AQZ10220.1); *Wickerhamiella_sorbophila* (ID: XP_024663738.1); *Geotrichum_candidum* (ID: CDO55024.1). Triangles and stars indicate key conserved amino acids, respectively. (D) Multiple pairs of conserved salt bridges are formed at the inter-subunit interface of the VTC complex. R83 of Vtc1(γ) and E704 of Vtc3 are separated by 8 Å, too far to form a salt bridge。 (E) Cellular polyP content of Vtc4p, Vtc3p and Vtc1p point mutants expressed under the control of their native promoters in the *vtc4*Δ, *vtc3*Δ(*vtc2*Δ) and *vtc1*Δ backgrounds, respectively. Δ indicates that the entire subunit was knocked out. FL indicates full length, indicating that the subunit has not been modified in any way. Data show the mean±s.d (n=3). (F) A model of the putative polyP channel gating mechanism. The schematic drawing illustrates the polyP channel conformational changes between the closed and open. The number 1 represents the TM1 of each subunit of the VTC complex. (G) A model of the activation mechanism of the VTC complex. Schematic of the Vtc4/3/1 complex. Subunits are colored. The three subunits of Vtc1 are shown in grey. Key amino acids are highlighted. The stars represent the binding sites of PP- InsPs or ATP.

Furthermore, we observed an unusual salt bridge at the center of the inter-subunit interface among the mainly hydrophobic interactions. E30 and R83 of Vtc1 are strictly conserved among Vtc1, Vtc2, Vtc3 and Vtc4 from different species (Figure 7C). R83 from the principal (+) Vtc1 subunit forms a salt bridge with E30 of the complementary (-) Vtc1 subunit. The corresponding residues are E704 and R762 of Vtc3, E628 and R681 of Vtc4 (Figure 7D). Similar salt bridges exist between Vtc3 and Vtc4, and Vtc4 and Vtc1(α). However, R83 of Vtc1(γ) and E704 of Vtc3 are separated by 8 Å, too far to form a salt bridge (Figure 7D). To confirm the importance of the observed inter-subunit salt bridge, we created 12-point mutants in the VTC complex and measured the cellular polyP content. All substitutions designed to disrupt the inter-subunit salt bridge by charge reversal or charge removal significantly reduced cellular polyP content (Figure 7E), indicating that the inter-subunit salt bridges are necessary for VTC complex function. To our surprise, substitution of E704 of Vtc3, which does not form inter-subunit salt bridge in the structure, also significantly reduced cellular polyP content (Figure 7E). We hence spectulate that E704 of Vtc3 might form such inter-subunit salt bridge in another functional state of VTC, for example during VTC channel opening and transit of a polyP chain. Due to their role in VTC complex function we term these salt bridges as inter-subunit “ionic locks”.

The flexible nature of the inter-subunit interface, together with the observation of inter-subunit “ionic locks”, suggests a plausible polyP channel gating mechanism. We assume that the polyP channel was captured in a resting state with the entrance fastened by a latch-like, horizonal helix of Vtc4. The asymmetrical nestling of the horizonal helix of Vtc4 at the entrance of the channel imposes forces asymmetrically, resulting in different relative positioning at the inter-subunit interfaces. Three loosely packing inter-subunit interfaces, together with two tightly packing ones, render the TM1 helices tapering from the cytosolic side towards the intravacuolar side, generating a narrow point that might serve as a gate. The inter-subunit “ionic locks” may hold the subunits together. During opening of the polyP channel, the horizontal helix latch is lifted, and the subunits are pulled together by the “ionic locks”, possibly resulting in the formation of all five “ionic locks” between subunits. In this state, the TM1 helices might tilt inwards at the cytosolic side and outwards at the intravacuolar side, thus opening the channel (Figure 7F).

## Discussion

### The observed central domain of Vtc4 exists in a polyP polymerase “off” state

Three lines of evidence lead us to believe that the observed central domain of Vtc4 in our cryo-EM structure is in a polyP polymerase “off” state. First, polyP synthesis and the immediate translocation of polyP into the vacuole are coupled (Gerasimaite et al., 2014), and our structure of the Vtc4/Vtc3/Vtc1 complex supports such a coupled polyP polymerase and translocase. Given that the polyP channel in our structure of the Vtc4/Vtc3/Vtc1 complex was captured in a resting state, it is then reasonable to assume that the polymerase is in an “off” state. Second, without the constraint from the polyP channel, the central domain of Vtc4 alone can synthesize polyP (Hothorn et al., 2009), indicating that the crystal structure of the central domain of Vtc4 represents a polymerase “on” state. Indeed, the electron density for polyP had been observed in one of the crystal structures of the central domain of Vtc4 (Hothorn et al., 2009). While no polyP product was observed in the cryo-EM structure of the whole Vtc4/Vtc3/Vtc1 complex. Third, structural comparison revealed that the β4-β5 loop of the central domain of Vtc4 adopts a different conformation between the crystal structure of the central domain of Vtc4 alone (Hothorn et al., 2009) and the cryo-EM structure of the whole Vtc4/Vtc3/Vtc1 complex (Figure S16). Indeed, the β4-β5 loop of the central domain of Vtc4 is very sensitive to mutagenesis. Substitutions in this region abolished polyP synthesis in vitro (E290G, E290A, E290R) and in vivo (R284A/E285A/D286A) (Figure S16B, S16C).

### The activation mechanism of the VTC complex

The data presented here have allowed us to elucidate a possible activation mechanism of the VTC complex. We suppose that the VTC complex exists in equilibrium between an inactive and active state. ATP and the inositol-based signaling molecules activate the VTC complex. The Vtc4/Vtc3/Vtc1 structure was captured in an inactive state, with a resting state polyP channel and an “off” state polyP polymerase. How might the VTC complex convert the free energy of ATP binding, or the binding of the inositol-based signaling molecules to turn “on” the polyP polymerase, and open the polyP channel? To address this question, we superimposed the structures of isolated SPX domains (SPX^CtGde1^-InsP6 (PDB ID: 5IJJ), or the SPX^CtVtc4^-InsP6 (PDB ID: 5IJP)) (Wild et al., 2016) to the SPX domain of Vtc4 of the intact VTC complex, and observed that the InsP6 bound on the SPX domain is close to Vtc3 (Figure S17). The SPX domain harbors a large, positively charged surface able to interact with phosphate-containing inositol ligands but showing little specificity and selectivity at the level of binding (Wild et al., 2016). One can imagine a phosphate-containing ligand, such as ATP, 5-IP7 or 1,5-IP8, binds in the cleft between the large, positively charged surface of the SPX domain of Vtc4 and the α1 helix of the central domain of Vtc3 and causes the domains to move relative to each other, thus inducing conformational change that turns “on” the polyP polymerase, followed by the opening of the polyP channel. In addition, it is worth noting that the binding affinity of phosphate-containing ligands to SPX domain, gradually increase from P_i_, pyrophosphate (PP_i_), triphosphate (PPP_i_), to InsP6, with a 20-fold higher K_d_ value of InsP6 than that of PPP_i_ (Wild et al., 2016). This allowed us to propose a simplified model of the activation and regulation in VTC complex (Figure 7G). The VTC complex contains a polyP polymerase, a polyP channel and a regulatory cleft, and exists in equilibrium between inactive or active states. The high apparent Km value of VTC for ATP might be an additional mechanism to reduce VTC activity in situations where Pi is abundant but the cells cannot generate sufficient ATP. Then, the high Km would provide an inbuilt mechanism to reduce polyP synthesis, which may be a strong consumer of ATP but dispensable for survival under such conditions. PP-InsPs might serve as high affinity stimulatory ligands when ATP and Pi are abundant. Also the synthesis of InsPPs itself is probably impacted by the ATP concentration, because both the IP6 kinases and PPIP kinases, which synthesize IP7 and IP8, have high Km values for ATP (Nair, Gu et al., 2018, Voglmaier, Bembenek et al., 1996), which are close to the ATP concentrations in the cytosol. Thus, the VTC complex may integrate information about the ATP and phosphate status of the cell at two levels. Such control at multiple levels may be justified by the fact that VTC is a powerful phosphate pump, which can push the cells into phosphate starvation when improperly regulated (Austin & Mayer, 2020, Desfougeres et al., 2016).

## Materials and Methods

### Yeast strains and plasmids

The protease-deficient *S. cerevisiae* BJ2168 (MATa: leu2-3, trp1-289, ura3-52, prb1-1122, pep4-3, prc1-407, gal2) was used as a host strain. The modified TAP tag (6His-TEV-Protein A, named TAPm tag) or the strep tag was inserted at the C-terminus of Vtc2 or Vtc3 by a homologous recombination-based method (Funakoshi & Hochstrasser, 2009). Based on the above methods, we constructed Vtc2-TAPm, Vtc3-TAPm and Vtc3(ΔC24)-TAPm single-tag strains, as well as Vtc2-TAPm/Vtc3- Strep and Vtc2-Strep/Vtc3-TAPm dual-tag strains.

Single subunit knockout strains Vtc1Δ, Vtc2Δ, Vtc3Δ and Vtc4Δ were prepared using a plasmid pYM27-kanMX in the BJ2168 strain. The kanMX gene replaces the VTC gene behind the promoter of the corresponding subunit. Double subunits knockout strain Vtc2Δ and Vtc3Δ were prepared using a plasmid p426-URA3 in the Vtc3Δ strain.

The genes of Vtc1, Vtc2, Vtc3 and Vtc4 were cloned into plasmid p426-URA3 for various site-directed mutagenesis. Vtc1 point mutants were expressed from p426- URA3 plasmid integrated into the genome behind the Vtc1 promotor of a VTC1::kanMX strain. Similarly, Vtc2, Vtc3 and Vtc4 point mutants were performed in the same way.

*S. cerevisiae* Ppx1 was cloned into pET28a (kanamycin (Kan) selection) vector and transferred to BL21 (DE3) for expression.

### Purification of the endogenous wild type and mutant VTC complexes

Yeast cells were cultured in YPD (2% peptone, 1% yeast extract, 2% glucose) medium for 18 h. The cells were collected by centrifugation at 4,000 rpm at 4°C. The collected cells were resuspended in lysis buffer containing 25 mM Hepes-NaOH (pH 7.4), 150 mM NaCl, 1 mM EDTA, and supplemented with a protease inhibitor cocktail (2 μg/ml DNase I, 1 μg/ml pepstatin, 1 μg/ml leupeptin and 1 μg/ml aprotinin, and 1 mM PMSF) and then were lysed using a high pressure homogenizer at 1,000 bar for 5 cycles. After lysis, cell debris was pelleted by centrifugation at 8,000g for 10 min. The supernatant was subjected to centrifugation in a Ti45 rotor (Beckman) at 40,000 r.p.m. for 1 h at 4°C. The pelleted membranes were resuspended with a Dounce homogenizer in buffer A containing 25 mM Hepes-NaOH (pH 7.4), 150 mM NaCl, 1 mM EDTA, 1 mM MgCl_2_, 1 mM MnCl_2_, 1 mM PMSF, and 2% DDM + 0.2% CHS. After incubation for 3h at 4 °C, the mixture was centrifuged for 30 min at 18,000 r.p.m to remove insolubilized membrane. The supernatant was incubated with 600 μl IgG resin for 3 h. The beads were washed with 30 ml buffer B (25 mM Hepes-NaOH (pH 7.4), 150 mM NaCl, 1 mM MgCl_2_, 1 mM MnCl_2_, 0.1% digitonin), and the complex was eluted with buffer B containing 0.15 mg/ml TEV protease. The complex was concentrated and further purified by size-exclusion chromatography on a Superose 6 10/300 Increase column, equilibrated with buffer B. Peak fractions were pooled and concentrated to 8 mg/ml for cryo-EM analysis.

### Cryo-EM grid preparation and data collection

For the cryo-EM grids preparation, 3ul purified Vtc4/Vtc3/Vtc1 complex at a concentration of about 8 mg/ml was applied respectively to glow-discharged holey carbon EM grids (CryoMatrix Amorphous alloy film R1.2/1.3, 300 mesh). The grids were blotted for 3 s with a blot force of 0 and then plunged into liquid ethane using a FEI Vitrobot Mark IV (Thermo Fisher Scientific) at 4°C and 100% humidity. The cryo-EM grids were subsequently transferred into a 300kV Titan Krios microscope (Thermo Fisher Scientific) equipped with a Gatan K3 direct electron detector and a BioQuantum energy filter operated at a slit width of 20 eV. Micrographs were automatically collected by EPU in super-resolution mode with a pixel size of 0.4255Å. Each micrograph was comprised of 40 frames with a total exposure time of 2.5s and total dose of 54 electrons per Å^2^. The defocus range for each micrograph was set from -1.0 to -1.5μm.

### Cryo-EM data processing

The collected movie stacks were summed and corrected for beam-induced motion using MotionCor2 (Zheng, Palovcak et al., 2017) with a binning factor of 2. Gctf (Zhang, 2016) was used for estimating contrast transfer function (CTF) parameters for each micrograph. And the following processing steps including particle picking, 2D classification, 3D classification, 3D auto-refine, CTF refinement and polishing were all performed using RELION-3.1.1 (Zivanov, Nakane et al., 2020). Local resolution map was estimated using RELION. All 3D density maps were displayed using UCSF Chimera (Pettersen, Goddard et al., 2004).

For the Vtc4/Vtc3/Vtc1 complex, 3871 and 3493 micrographs were collected separately. And a total of 1,641,408 and 1,764,870 particles were auto-picked and subjected to 2D classification and 3D classification individually. After that, good classes showed clear features were combined from two datasets including 1,542,410 particles and subjected to another round of 3D classification. And two best-resolved classes were chosen and combined containing 1,042,873 particles for 3D refinement, CTF refinement and polishing. The final refinement generated a map with a global resolution of 3.06 Å. And signal subtract was used for a more detailed feature and higher resolution map of Transmembrane region.

### Cryo-EM model building, refinement and validation

The initial templates of Vtc1, Vtc3 and Vtc4 were generated using AlphaFold2 (Jumper, Evans et al., 2021). The transmembrane domain, similar catalytic domain and SPX domain of Vtc3 and transmembrane domain, catalytic domain and SPX domain of Vtc4 were cut out and separately rigid body fitted into cryo-EM density map using Chimera (Pettersen et al., 2004). Then three copies of Vtc1 were docked into the remaining density map. The initial fitting of Vtc4/Vtc3/Vtc1 complex was confirmed by high agreement of secondary structural features between the predicted 3D models and the cryo-EM density map. Polyphosphate and POV coordinates and geometry restraints were generated using a phenix.elbow (Adams, Afonine et al., 2010) and fitted into density map. All the models were manual adjusted and rebuild using Coot (Emsley & Cowtan, 2004), followed by several round of real-space refinement in PHENIX (Adams et al., 2010) and manual adjustment in COOT (Emsley & Cowtan, 2004). The final models statistics were validated and provided by MolProbity (Williams, Headd et al., 2018) and summarized in Table S1. Structural figures were prepared using Chimera (Pettersen et al., 2004).

### Purification of recombinant ScPpx1

*E.coli* BL21(DE3) cells were grown in LB medium containing 50 μg/ml Kan at 37°C. 0.4 mM IPTG was added when OD600 reached 0.6. The cells were transferred to 16 °C and cultured for 18h before harvesting. Cell pellets were resuspended in lysis buffer containing 50 mM Hepes-NaOH (pH 7.4), 300 mM NaCl and disrupted by sonication. After lysis, cell debris was removed by centrifugation at 18,000 r.p.m for 30 min. The supernatant was incubated with 2 ml Ni-NTA resin for 30 min. The beads were washed with 30 ml lysis buffer plus 20 mM imidazole, followed by a second wash with 30 mL of lysis buffer plus 50 mM imidazole. The protein was eluted with lysis buffer plus 250 mM imidazole. The eluted protein was dialyzed against 50 mM Hepes-NaOH (pH 7.4), 150 mM NaCl to remove imidazole. Dialyzed protein was concentrated using an Amicon Ultra concentrator (30 kDa MWCO, Millipore) and aliquoted into 100 μl amounts and stored at −80 °C.

### Detection of PolyP content *in vivo*

Yeast cells (including wild-type strains, knockout strains and mutant strains) were grown overnight in YPD medium. Dilute all yeast cultures to an OD600 value of 1, 2 ml of the culture was centrifuged at 4000 rpm for 5 min, and cells were collected.

PolyP extraction and purification are based on an improved method (Bru, Jimenez et al., 2016). The cell pellet was resuspended with 400 μl of AE buffer (50 mM sodium acetate (pH 5.3), 10 mM EDTA) and add 300 μl phenol and 40 μl 10% SDS, mixed by inversion 4 times, vortexed 5 sec to homogenize, incubated at 65°C for 10 min and chilled for 2 min on ice. 300 μl of chloroform were added, mixed by inversion 4 times, vortexed 5 sec to homogenize and centrifuged at room temperature for 2 min at 14,000 r.p.m. The top aqueous phase was transferred to a prepared 1.5 ml screw cap tube containing 350 μl chloroform, mixed by inversion 4 times, vortexed 5 sec to homogenize, centrifuged at room temperature for 2 min at 14,000 r.p.m and the aqueous phase was transferred to a new 1.5 ml microcentrifuge tube. 2 μl of RNAse A(10 mg/ml) and 2 μl of DNAse I(10 mg/ml) were added, incubated 1 h at 37°C, transferred to a pre-cold at -20°C 1.5 ml microcentrifuge tube containing 1 ml of absolute ethanol and 40 μl of 3 M sodium acetate (pH 5.3), leaved 3 h at -20°C to precipitate polyP and centrifuged for 20 min at 14,000 r.p.m at 4°C. The supernatant was discarded, 500 μl of 70% ethanol were added, centrifuged for 10 min at 14,000 r.p.m at 4°C, the supernatant was discarded. The tube was left open to dry the small translucent-white polyP pellet at room temperature for 10 min or until the pellet is completely dry. Finally, the polyP was resuspended in 50 μl of deionized water. The polyP sample can be directly measured or stored at -20°C.

The purified polyP samples were measured by Malachite Green Phosphate Assay Kits (Sigma, POMG-25H). First, PolyP needs to be degraded into Pi by the polyphosphatase Ppx1. A 50 μl reaction system containing 5 μl PolyP, 0.5 μg Ppx1 and reaction buffer (50 mM Hepes-NaOH, pH 7.4, 150 mM NaCl) was reacted at 37 °C for 1 h. The Malachite Green Phosphate Assay Kit is based on quantification of the green complex formed between Malachite Green, molybdate and free orthophosphate. The rapid color formation from the reaction can be conveniently measured on a spectrophotometer (600 - 660 nm). Standard phosphate was used for assay calibration.

### Detection of PolyP synthesis *in vitro*

PolyP synthesis was assayed in 15 μl samples consisting of reaction buffer (25 mM Hepes-NaOH (pH 7.4), 150 mM NaCl, 1 mM MgCl_2_, 1 mM MnCl_2_, 0.1% digitonin), 5 mM ATP and 6 μg purified endogenous proteins (Vtc4/2/1 complex, and Vtc4/3/1 complex). After the entire reaction was maintained at 4°C for 1 h, stop buffer (25 mM Hepes-NaOH (pH 7.4), 150 mM NaCl, 0.1% digitonin, 15 mM EDTA, 15 μM DAPI)was added until the total volume reached 200 μl. The addition of EDTA chelated metal ions and destroyed the catalytic activity of the VTC complex. DAPI can form a complex with synthetic polyP, so we were able to measure the production of PolyP based on the characteristic fluorescence emission of DAPI-polyP complex at 550nm. A total of 200 μl of the sample was transferred into a black 96-well plate and fluorescence was measured with a SPECTRAmax GEMINI XS fluorescence plate reader (Molecular Devices) using λex=415 nm, λem=550 nm at 27°C (Gerasimaite et al., 2014).

### InsP6 stimulates the synthesis of PolyP *in vitro*

PolyP synthesis was assayed in 15 μl samples consisting of reaction buffer (25 mM Hepes-NaOH (pH 7.4), 150 mM NaCl, 1 mM MgCl_2_, 1 mM MnCl_2_, 0.1% digitonin), 5 mM ATP, 6 μg purified endogenous proteins (Vtc4/2/1 complex, and Vtc4/3/1 complex) and 10 mM InsP6. After the entire reaction was maintained at 4°C for 1 h, stop buffer (25 mM Hepes-NaOH (pH 7.4), 150 mM NaCl, 0.1% digitonin, 15 mM EDTA, 15 μM DAPI) was added until the total volume reached 200 μl. The addition of EDTA chelated metal ions and destroyed the catalytic activity of the VTC complex. DAPI can form a complex with synthetic PolyP, so we were able to measure the production of PolyP based on the characteristic fluorescence emission of DAPI-polyP complex at 550nm. A total of 200 μl of the sample was transferred into a black 96-well plate and fluorescence was measured with a SPECTRAmax GEMINI XS fluorescence plate reader (Molecular Devices) using λex=415 nm, λem=550 nm at 27°C.

### PolyP detection by PAGE gel

*In vivo* purified PolyP or *in vitro* synthesized PolyP was mixed with one volume of 2x TBE-Urea sample buffer (50% urea, 2x TBE, 0.25% xylene cyanol, 0.25% bromphenol blue). The sample was resolved electrophoretically using a 12% polyacrylamide gel (29:1 acrylamide /bis-acrylamide) containing 7 M urea in TBE buffer pH 8.3, at 250 V/h for 2.5 h at 4°C. The dimensions of the gel were 200 mm height, 200 mm wide and 1.5 mm thick. The gel was stained by soaking it in the staining solution (25% methanol, 5% glycerol, 2 µg/ml DAPI, 50 mM Tris pH 10.5) for 30 min, and destained by soaking it in destaining solution (25% methanol, 5% glycerol, 50 mM Tris pH 10.5) for 1 h. Finally, to visualize the polyP the gel was exposed to 254 nm UV light in Syngene G-BOX trans-illuminator.

### Western blot detection of the interaction between Vtc2 and Vtc3

The Vtc2-TAPm/Vtc3-Strep and Vtc2-Strep/Vtc3-TAPm strains were collected, followed by disruption, membrane solubilization with detergent, and centrifugation.

The supernatant was incubated with IgG beads for 2h, followed by washing, and the protein was eluted by TEV protease. Add reducing SDS sample buffer to the samples and incubate at 75°C for 5 min. Vtc2 and Vtc3 were detected using anti-His and anti-Strep antibodies.

### Isolation of vacuoles

The cells were grown in 1 litre of YPD medium at 30°C overnight and harvested at an OD_600_ of 0.6–1.3. A total of 600 ml of culture was centrifuged (2 min, 3900 g), cells were resuspended in 50 ml of 0.1 M Tris-HCl pH 8.9, 10 mM DTT, incubated for 7 min at 30°C in a water bath and collected by centrifugation. Cells were resuspended in 15 ml of spheroplasting buffer (50 mM potassium phosphate pH 7.5, 600 mM sorbitol in YPD with 0.2% glucose), 3000–4500 units of lyticase (Cabrera and Ungermann, 2008) were added and cells were incubated for 26 min at 30°C in a water bath. Spheroplasts were collected by centrifugation (3 min, 3400 g, 4°C) and gently resuspended in 15 ml of 15% Ficoll 400 in PS buffer (10 mM PIPES/KOH pH 6.8, 200 mM sorbitol). Spheroplasts were lysed by adding DEAE-dextran to a concentration of 7 mg/l and incubated (2 min, 0°C, then 2 min, 30°C). Samples were chilled, transferred into SW41 tubes and overlaid with 2.5 ml of 8% Ficoll 400, 3.5 ml of 4% Ficoll 400, and 1.5 ml of 0% Ficoll 400 (all in PS buffer). After centrifugation (150,000 g, 90 min, 4°C), vacuoles were harvested from the 0–4% interface. When isolating vacuoles from proteolytically competent strains, 1 mM PMSF and 16 protease inhibitor cocktail (16 PIC – 100 mM pefabloc SC, 100 ng/ml leupeptin, 50 mM O-phenanthroline and 500 ng/ml pepstatin A) were included in all buffers, starting from the spheroplasting step. Vacuole amounts were determined by protein content, using the Bradford assay with fatty-acid-free BSA as standard.

### PolyP synthesis by isolated vacuoles

PolyP synthesis was assayed in 100-ml samples consisting of reaction buffer (10 mM PIPES/KOH pH 6.8, 150 mM KCl, 0.5 mM MnCl_2_, 200 mM sorbitol) and ATP-regenerating system (ATP-RS – 1 mM ATP-MgCl_2_, 40 mM creatine phosphate and 0.25 mg/ml creatine kinase). The reactions were started by adding 2 mg of purified vacuoles, the samples were incubated at 27°C, followed by addition of 200 ml of stop solution (12 mM EDTA, 0.15% Triton X-100 and 15 mM DAPI) in dilution buffer (10 mM PIPES/KOH pH 6.8, 150 mM KCl, 200 mM sorbitol). This threefold dilution with EDTA-containing buffer did not only stop nucleotide hydrolysis but also resulted in faster development of DAPI–polyP fluorescence. Given that DAPI is membrane impermeable, dissolving the membranes with detergent was required in order to detect the entire polyP pool. A total of 240 ml of the sample was transferred into a black 96- well plate and fluorescence was measured with a SPECTRAmax GEMINI XS fluorescence plate reader (Molecular Devices) using λex=415 nm, λem=550 nm (cutoff=530 nm) at 27°C. Fluorescence was read every 1–2 min until the signal was stable. Experiments were repeated with at least three independent vacuole preparations. Values are presented as the mean±s.d.

## ACKNOWLEDGMENTS

This work was supported in part by Ministry of Science and Technology (2020YFA0908500 to S.Y. and 2020YFA0908400 to S.W.), the National Natural Science Foundation of China (31971127 to S.Y. and 31900930 to S.W), China Postdoctoral Science Foundation (2020M672434 to S.W.), and the Fundamental Research Funds for the Central Universities (to S.Y.).

## Author contributions

W.L. prepared the protein samples for cryo-EM and performed functional assays with the assistance from M.Z. and Q.C.. J.W., X.Y. and S.W. performed cryo-EM sample preparation, acquired cryo-EM data, data processing and analysis. H.Y and L.M helped with cryo-EM data collection. V.C. and A.M. performed polyP synthesis assay by isolated vacuoles. A.M. provided important insights, and helped with manuscript preparation. S.Y. and W.L. initiated the project, planned and analyzed experiments, supervised the research, and wrote the manuscript with input from all co-authors.

## Data Availability

The Structure coordinates and cryo-EM density maps have been deposited in the protein data bank under accession number XXXX and EMD-XXXXX.

---

**Figure S1.**
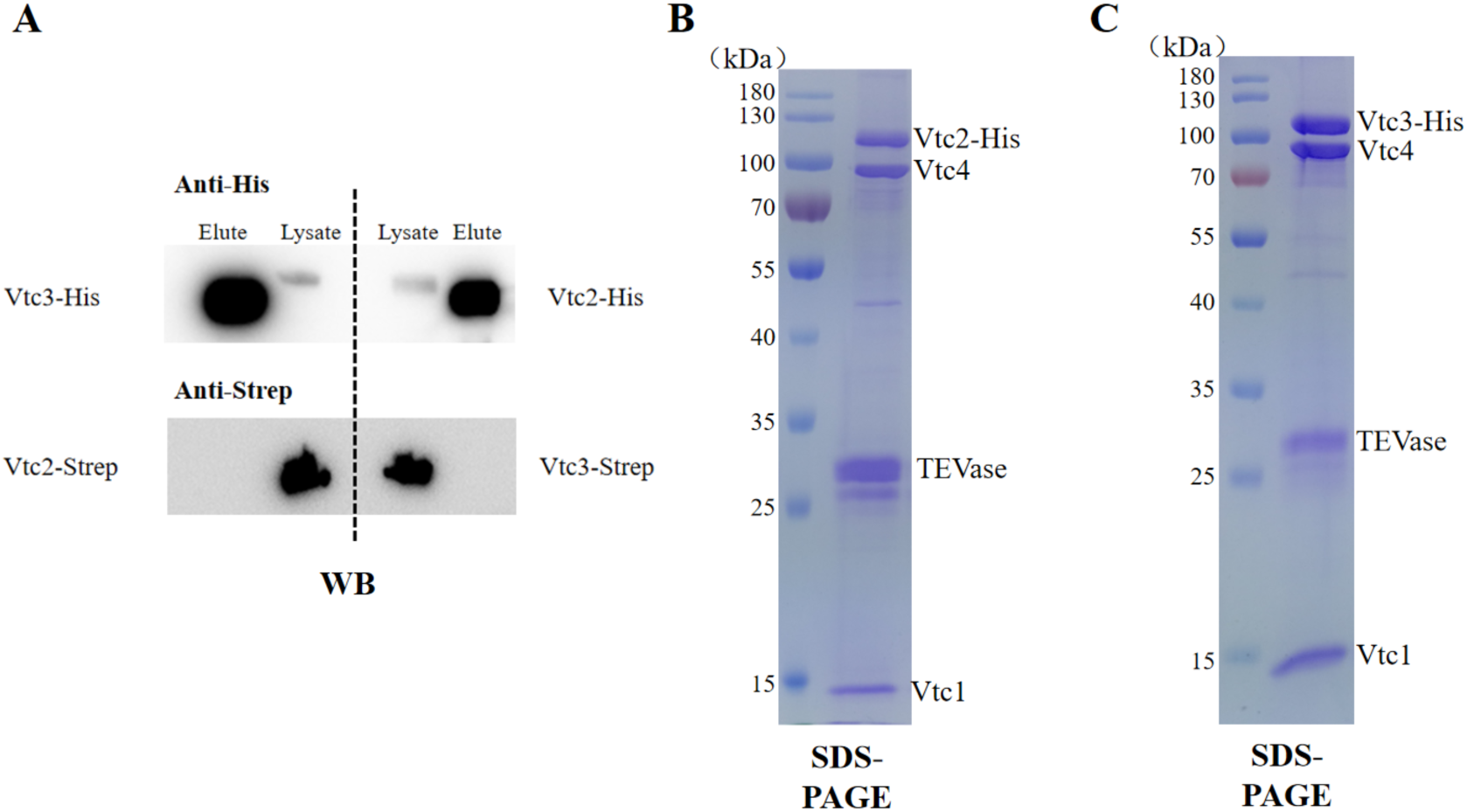
Purification of endogenous VTC complexes from *Saccharomyces cerevisiae*. (A) The interaction between Vtc2 and Vtc3 was detected by co-immunoprecipitation. The Vtc2-TAPm (-6His-TEV-Protein A)/Vtc3-Strep and Vtc2-Strep/Vtc3-TAPm strains were constructed. Whole cell lysate was incubated with IgG resin for 2h, followed by washing, and the protein was eluted by TEV protease. With the addition of reducing SDS sample buffer and incubation at 75°C for 5 min, the protein samples were run on SDS-PAGE gel. Vtc2 and Vtc3 were detected using anti- His and anti-Strep antibodies. (B and C) The Coomassie blue-stained SDS–PAGE gel of the purified (B) Vtc4/Vtc2/Vtc1 complex and (C) Vtc4/Vtc3/Vtc1 complex.

**Figure S2.**
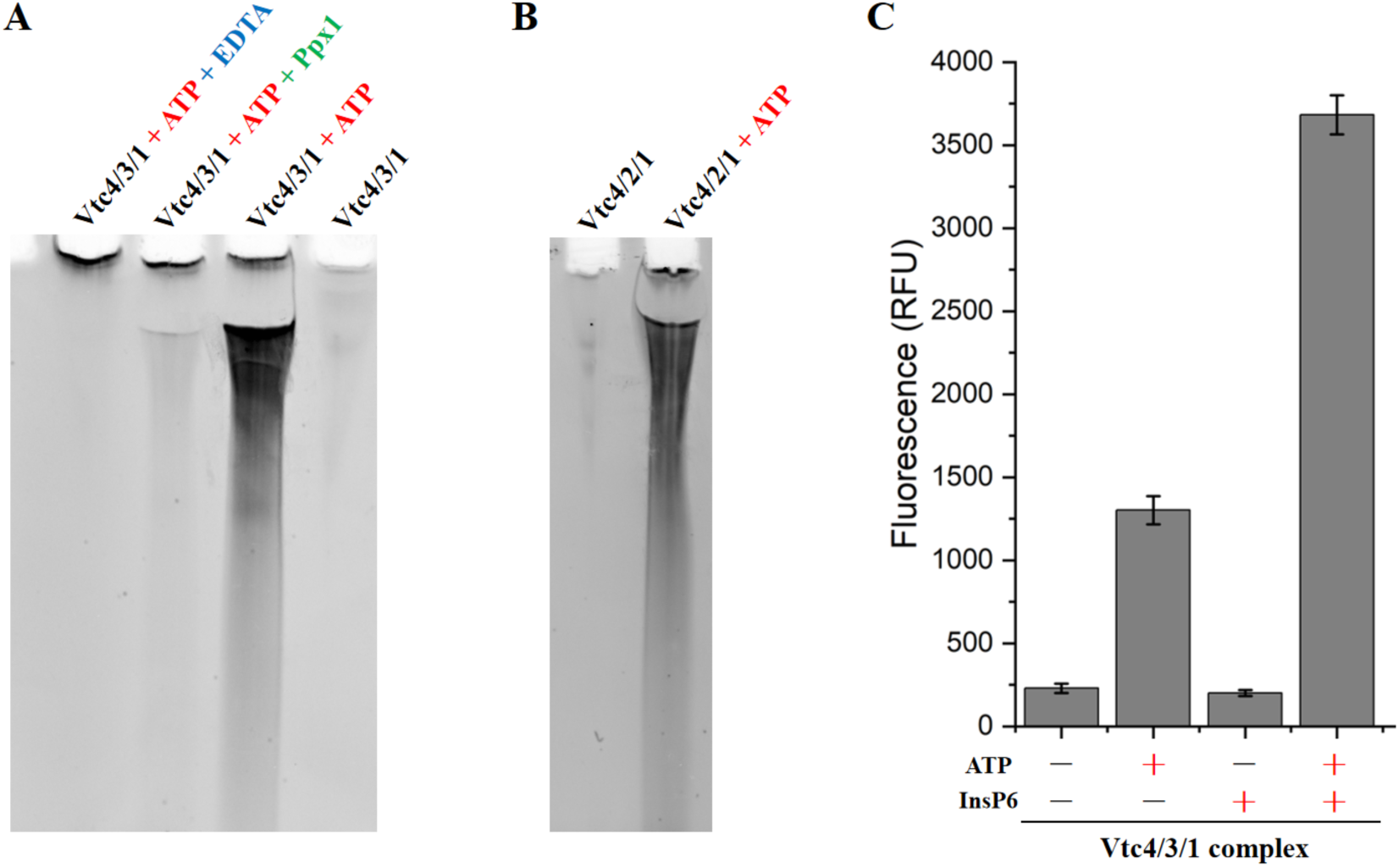
Purified endogenous VTC complexes synthesize polyP *in vitro*. (A and B) Urea-PAGE of polyP synthesized by (A) Vtc4/3/1 complex and (B) Vtc4/2/1 complex *in vitro*. Synthetic polyP was fractionated in a 12% polyacrylamide gel, and polyP was visualized by negative DAPI staining. Ppx1: a polyphosphatase from yeast that specifically hydrolyzes polyP. (C)The purified endogenous Vtc4/3/1 complex synthesizes polyP *in vitro* in an ATP- and InsP6-dependent manner. Data show the mean±s.d.

**Figure S3.**
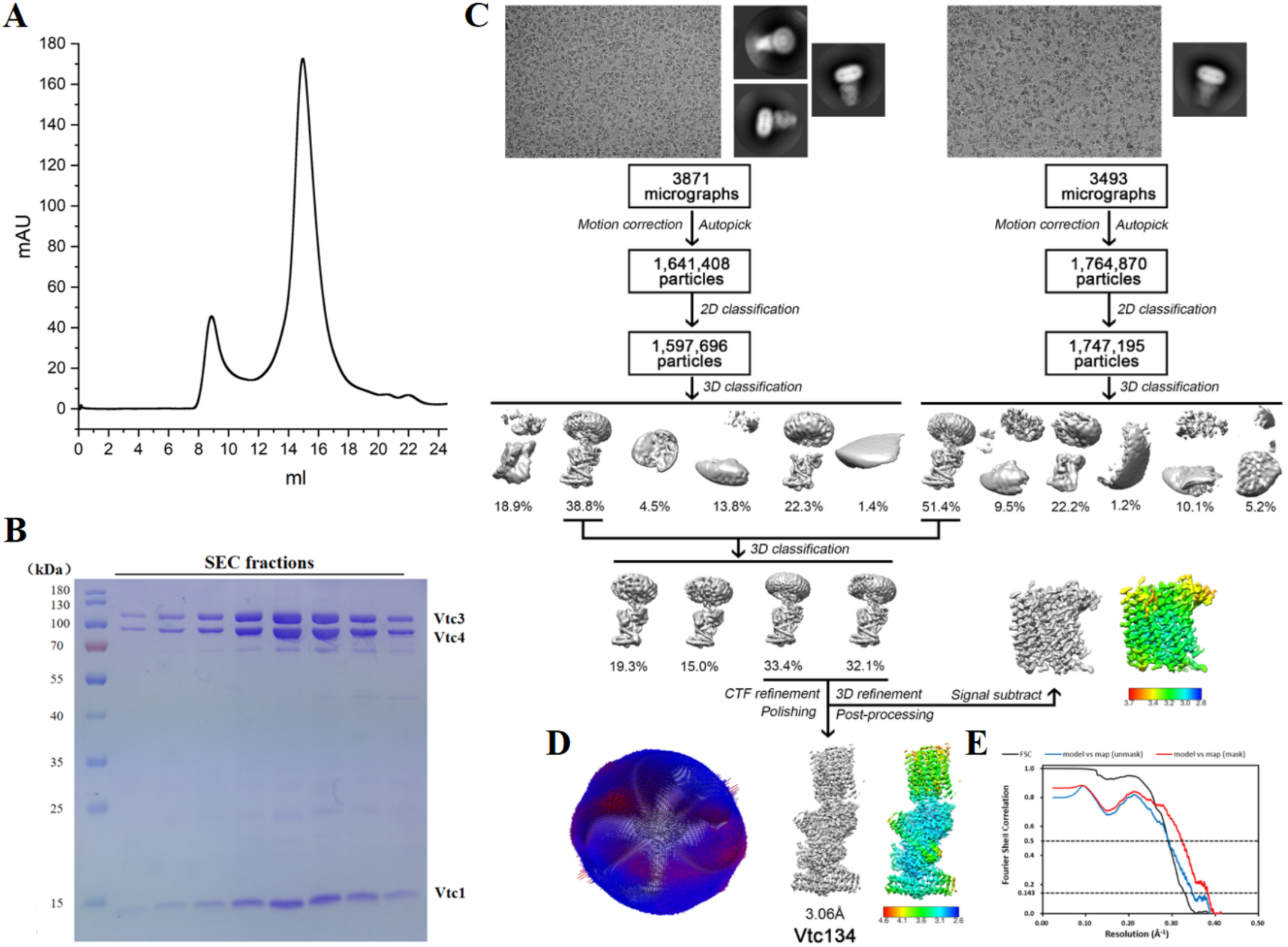
Cryo-EM image processing procedure of the Vtc4/Vtc3/Vtc1 complex. (A) Size-exclusion chromatography profile of the Vtc4/Vtc3/Vtc1 complex. (B) The Coomassie blue-stained SDS–PAGE gel of the pooled fractions from A. (C) Image processing workflow of the Vtc4/Vtc3/Vtc1 complex. (D) Angular distribution of particles used in the final reconstruction of the 3D map. (E) Gold-standard Fourier shell correlations of the final 3D reconstruction of the Vtc4/Vtc3/Vtc1 complex, and the validation correlation curves of the atomic model by comparing the model with the final map.

**Figure S4.**
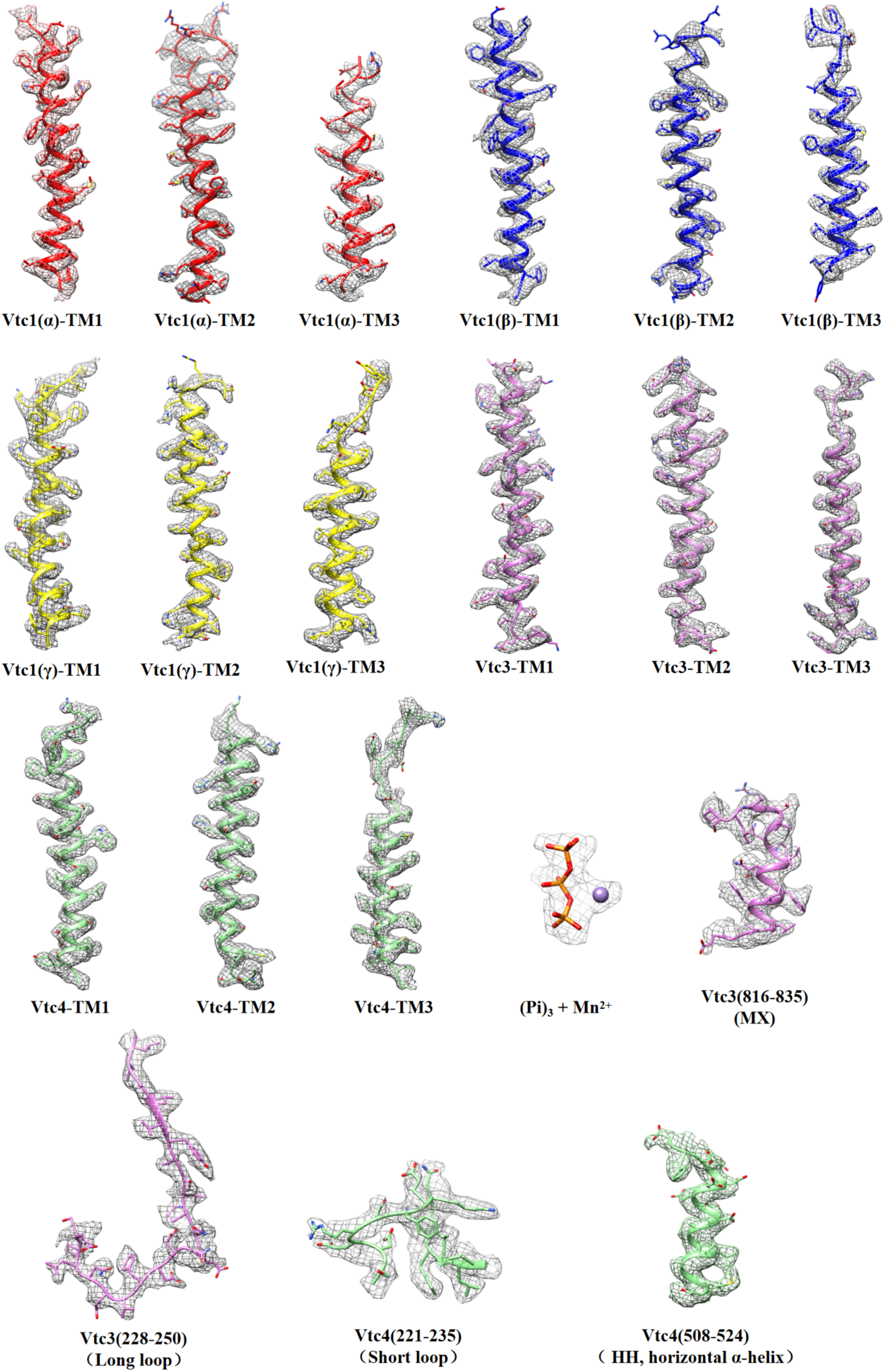
The fitting of the atomic model and the 3D map in selected regions. 3D density map and atomic model of selected regions in each of the five Vtc4/Vtc3/Vtc1 subunits, as well as the densities of atomic models of the triphosphate and Mn^2+^. MX, amphiphilic helix; HH, horizontal α-helix. For clarity, Vtc1(α), Vtc1(β) and Vtc1(γ) are colored in red, blue and yellow, respectively.

**Figure S5.**
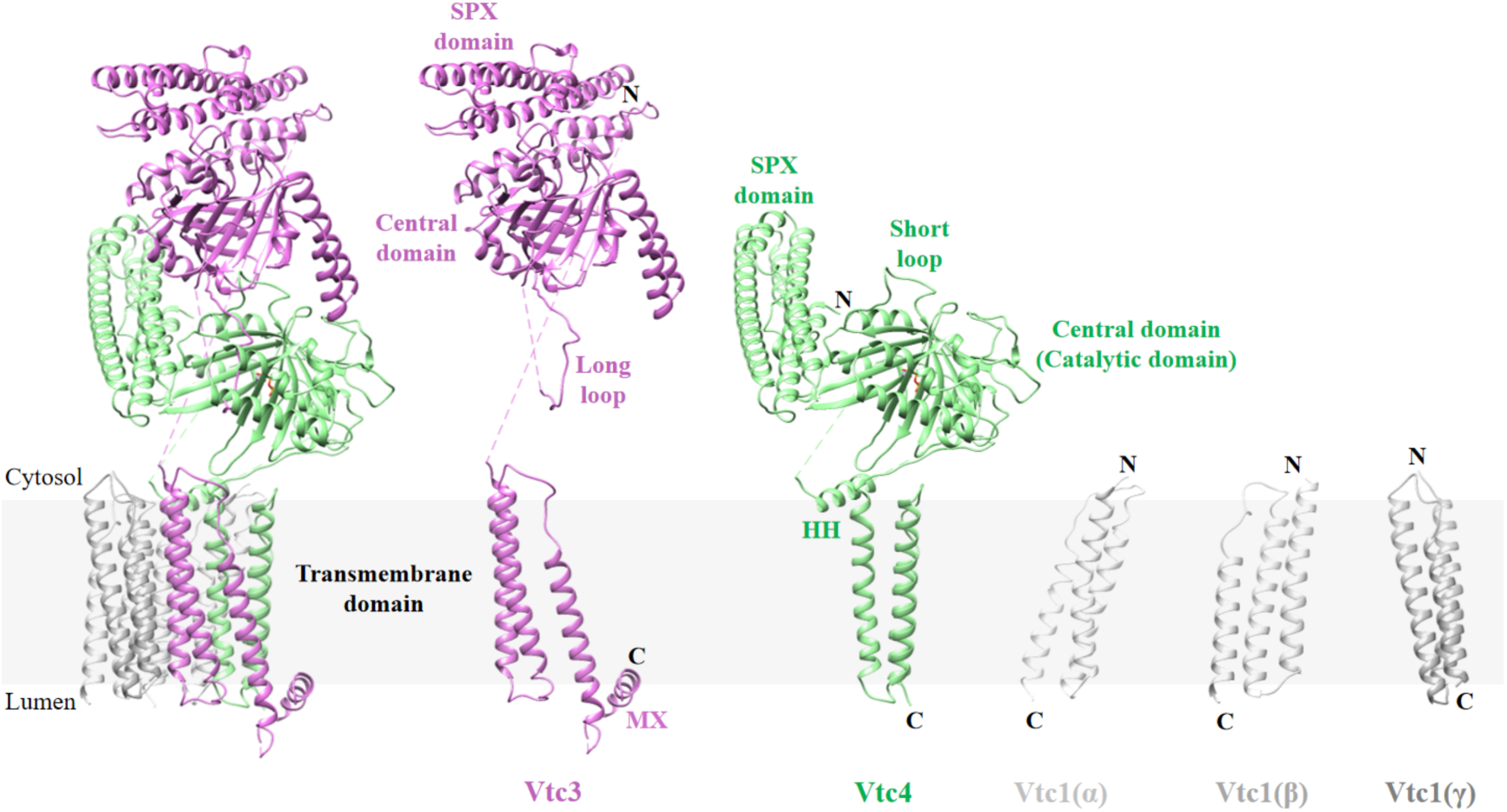
The structure of the five Vtc4/Vtc3/Vtc1 subunits. Structures of the five Vtc4/Vtc3/Vtc1 subunits shown separately. MX, amphipathic helix; HH, horizontal α-helix.

**Figure S6.**
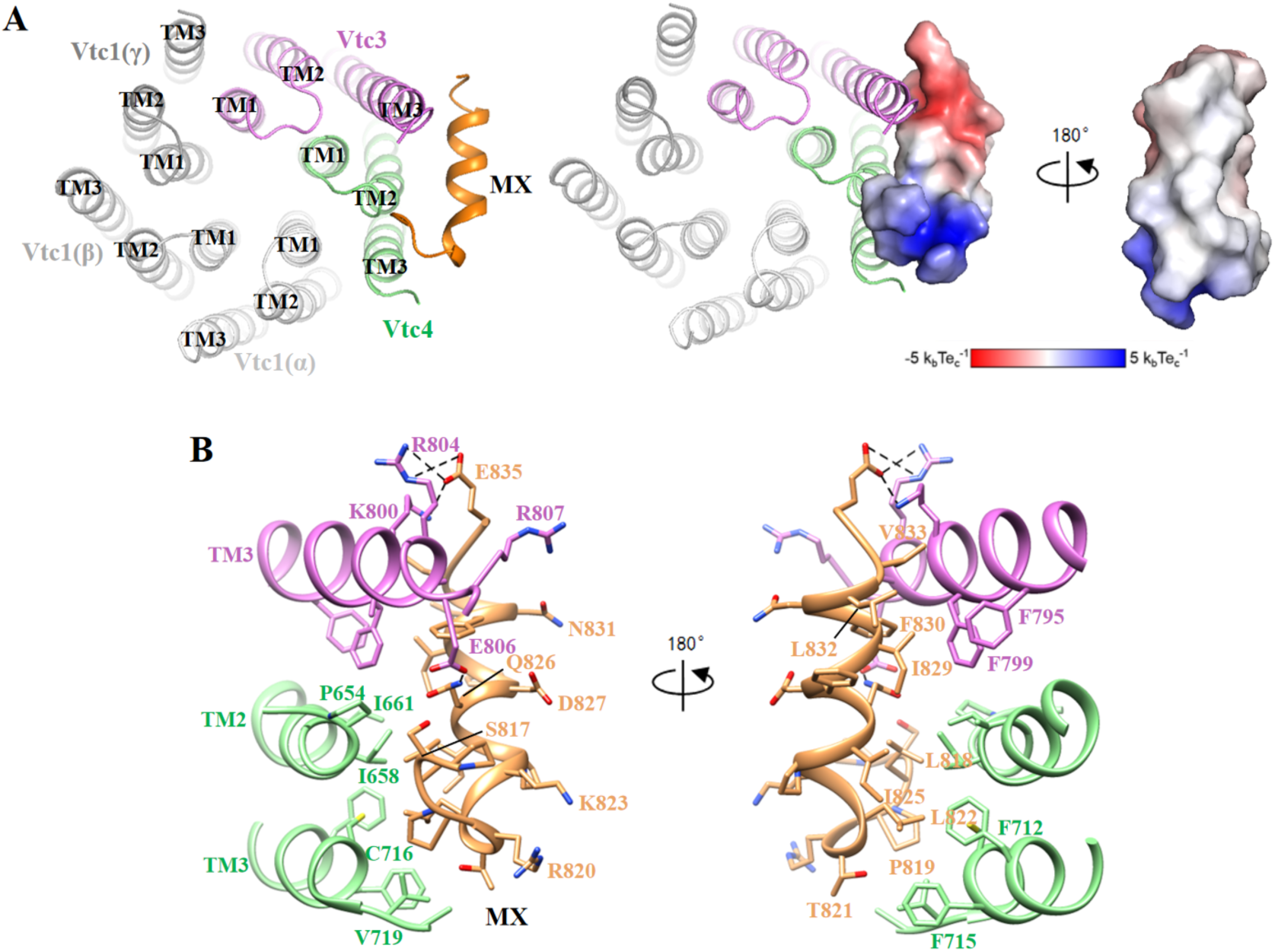
The structure of the MX helix of Vtc3. (A) Structure and electrostatic surface potential of the MX helix. The side of the MX helix close to the membrane is very hydrophobic, and the side away from the membrane is very hydrophilic. MX is shown in sandy brown. (B) The interaction between the MX helix of Vtc3 and the transmembrane domain.

**Figure S7.**
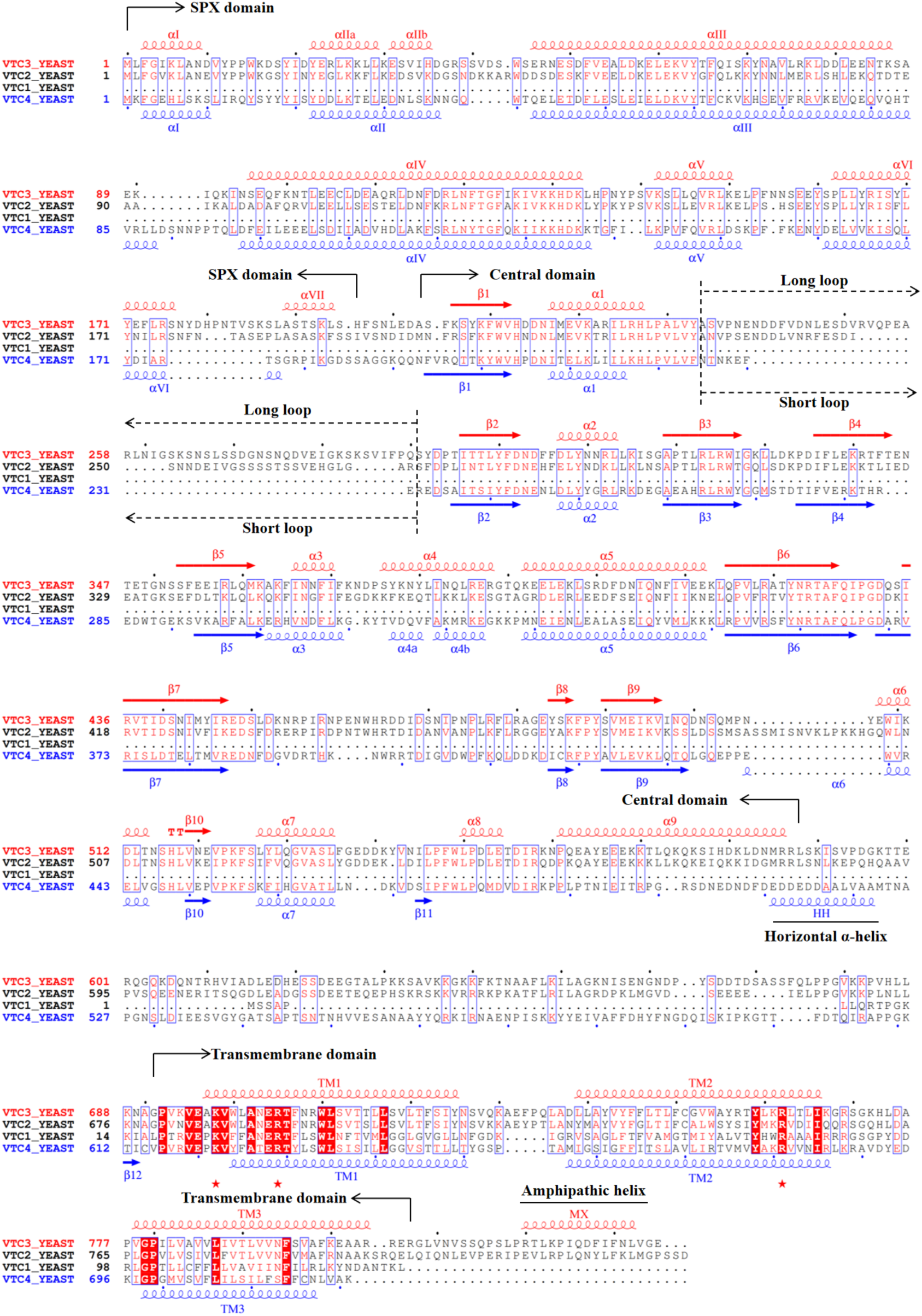
Sequence alignment of Vtc4, Vtc3, Vtc2 and Vtc1. Helices and β-strands are shown as coils and arrows, respectively. The assignments are produced by ESPript 3.0 (https://espript.ibcp.fr/ESPript/ESPript/) based on the structures S.c. Vtc3 (this study) and S.c. Vtc4 (this study). Key features are marked above the sequences. The red pentagram represents some conserved amino acids.

**Figure S8.**
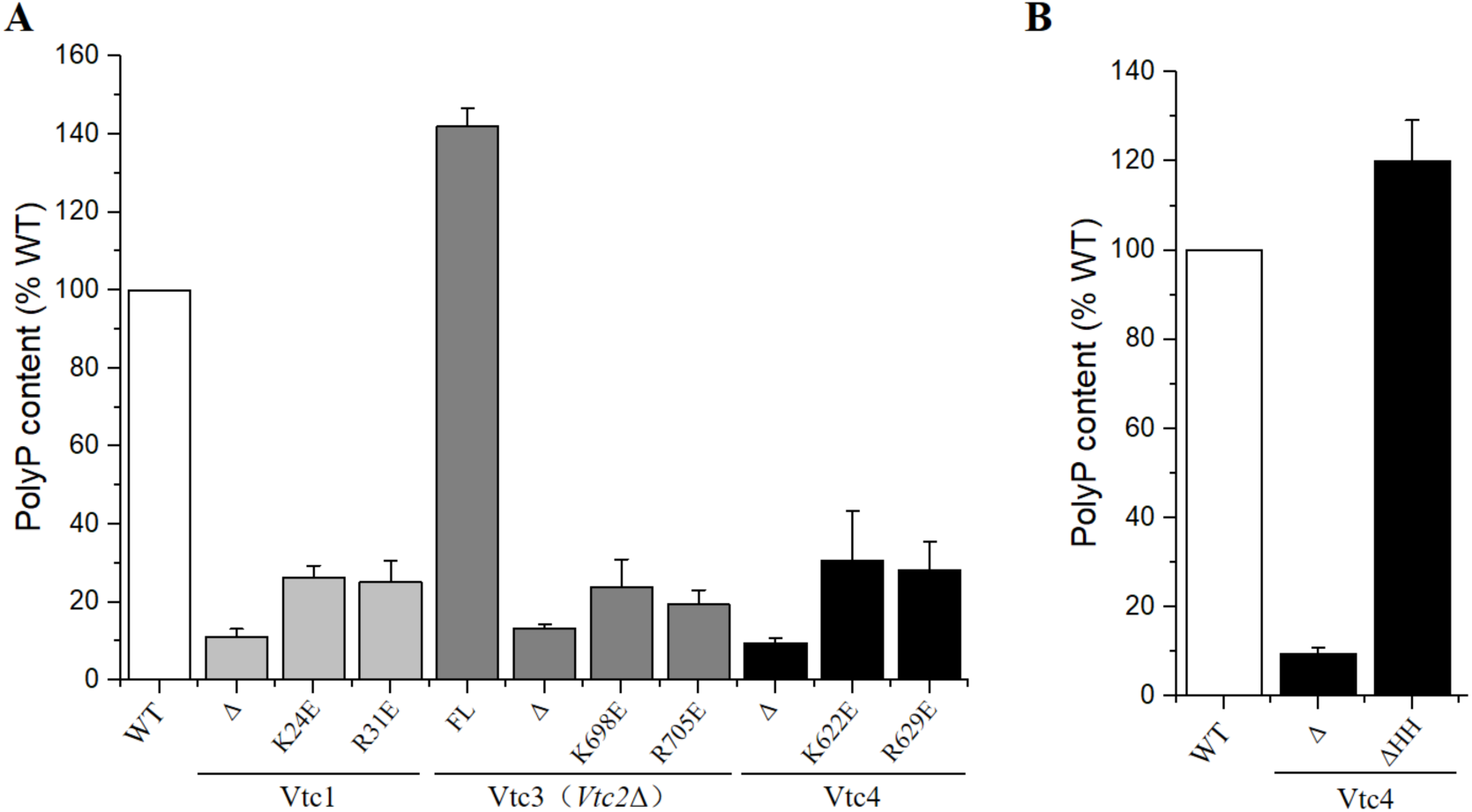
Mutational analysis of the polyP channel of the VTC complex. (A) Cellular polyP content of Vtc4p, Vtc3p and Vtc1p point mutants expressed under the control of their native promoters in the *vtc4Δ*, *vtc3Δ*(*vtc2Δ*) and *vtc1Δ* backgrounds, respectively. Δ indicates that the entire subunit was knocked out. FL indicates full length, indicating that the subunit has not been modified in any way. Data show the mean±s.d (n=3). (B) Cellular polyP content with truncated horizontal helices of Vtc4p expressed under the control of its native promoter in the *vtc4Δ* background. Δ indicates that the entire subunit was knocked out. FL indicates full length, indicating that the subunit has not been modified in any way. Data show the mean±s.d (n=3).

**Figure S9.**
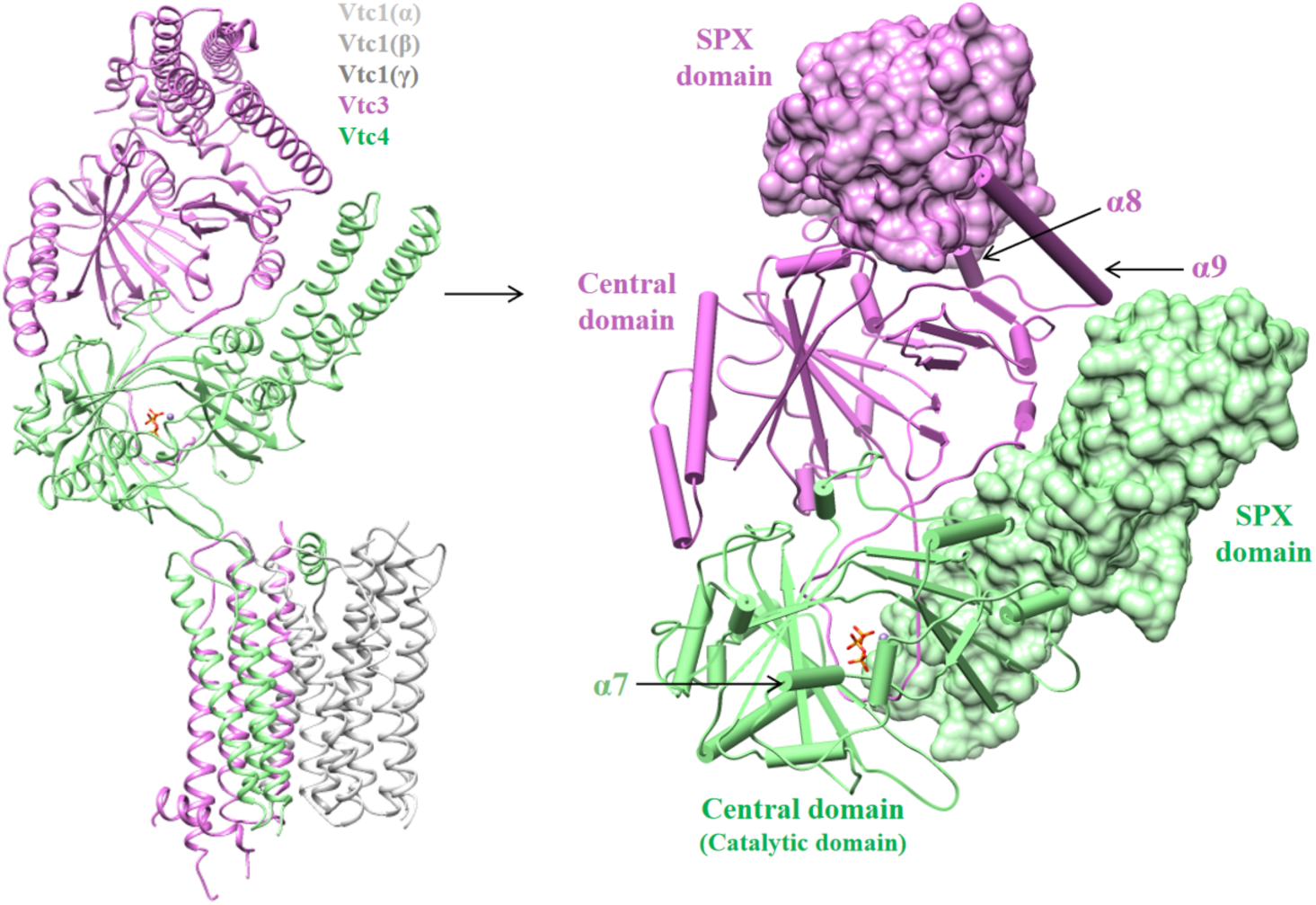
The cytoplasmic domain of Vtc3 and the cytoplasmic domain of Vtc4 form an asymmetric heterodimer. Both the cytoplasmic domain of Vtc3 and the cytoplasmic domain of Vtc4 contain an SPX domain and a central domain, and the central domain of Vtc3 has no catalytic activity.

**Figure S10.**
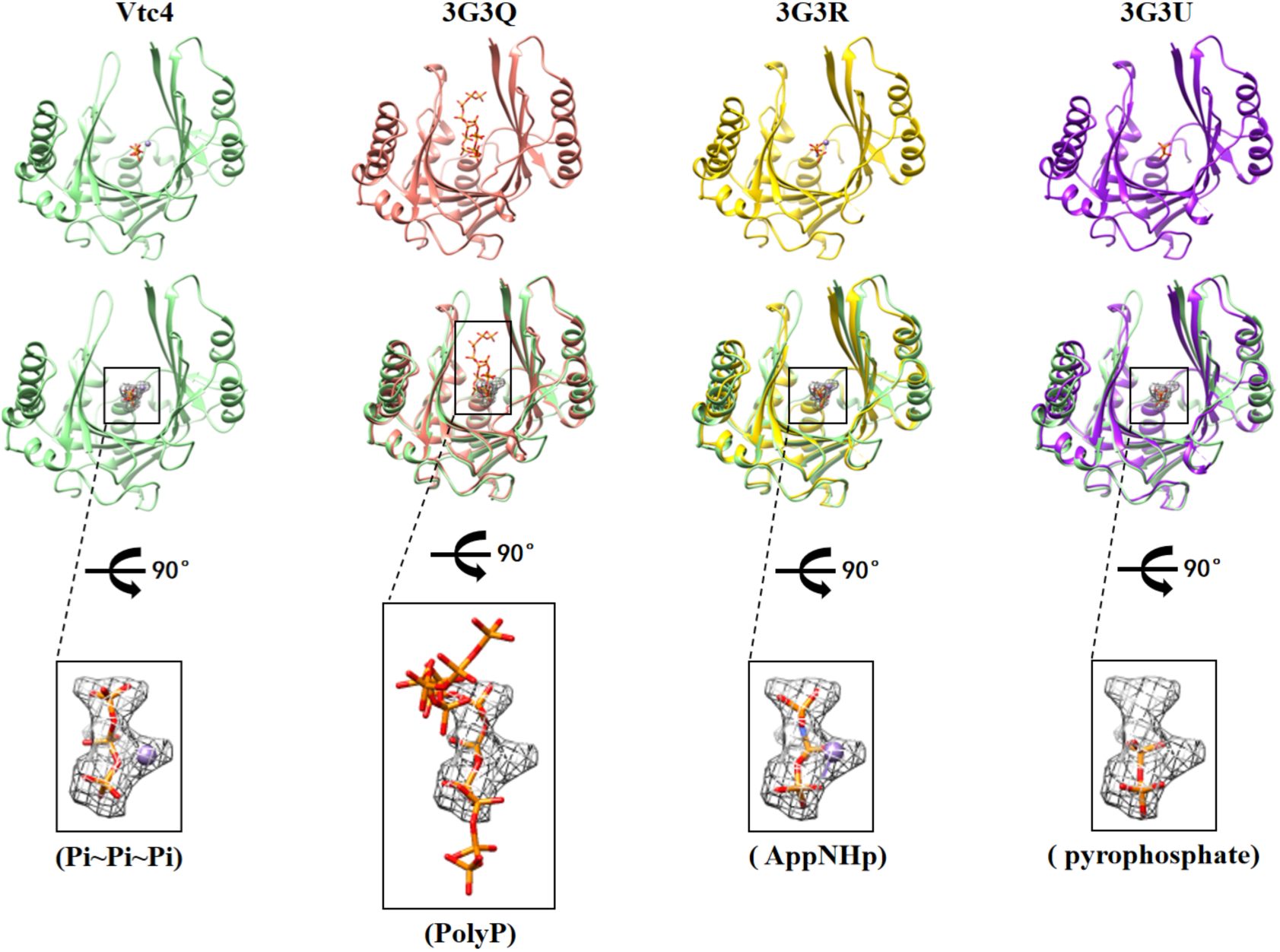
The structures of the central domain of Vtc4 (this study) were compared with those of the polyP-bound Vtc4 central domain (PDB: 3G3Q), the AppNHp-bound Vtc4 central domain (PDB: 3G3R) and the pyrophosphate- bound Vtc4 central domain (PDB: 3G3U), respectively. The positions of PolyP, AppNHp (adenosine-5’-[(β, γ)-imido] triphosphate) and pyrophosphate overlap with the triphosphates in our structure.

**Figure S11.**
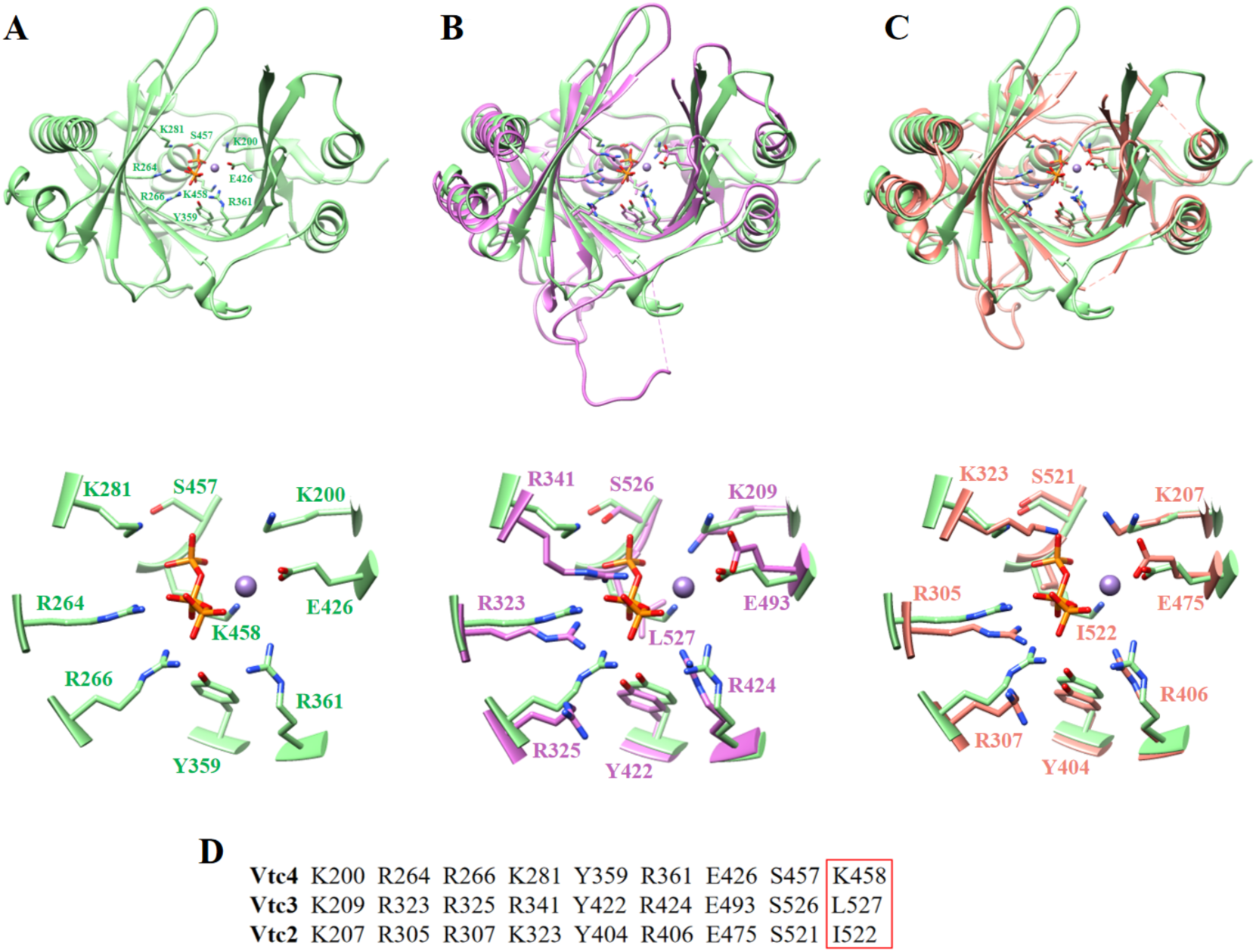
Structural comparison of the central domains of Vtc4, Vtc3 and Vtc2. (A) Structure of the central domain of Vtc4. The triphosphate and Mn^2+^ are shown in orange and brown, respectively. Some key residues are shown. (B) Compare the structures of the central domains of Vtc4 and Vtc3. The structures of the central domain of Vtc4 and Vtc3 are from our study. The key amino acids are highly conserved, only K458 of Vtc4 is replaced by L527 of Vtc3. (C) Compare the structures of the central domains of Vtc4 and Vtc2. The structure of the central domain of Vtc2 was obtained from the previously obtained crystal structure (PDB: 3G3O). The key amino acids are highly conserved, only K458 of Vtc4 is replaced by I522 of Vtc2. (D) Highly conserved amino acids in the central domain of Vtc4, Vtc3 and Vtc2. Non-conserved amino acids are circled in red boxes.

**Figure S12.**
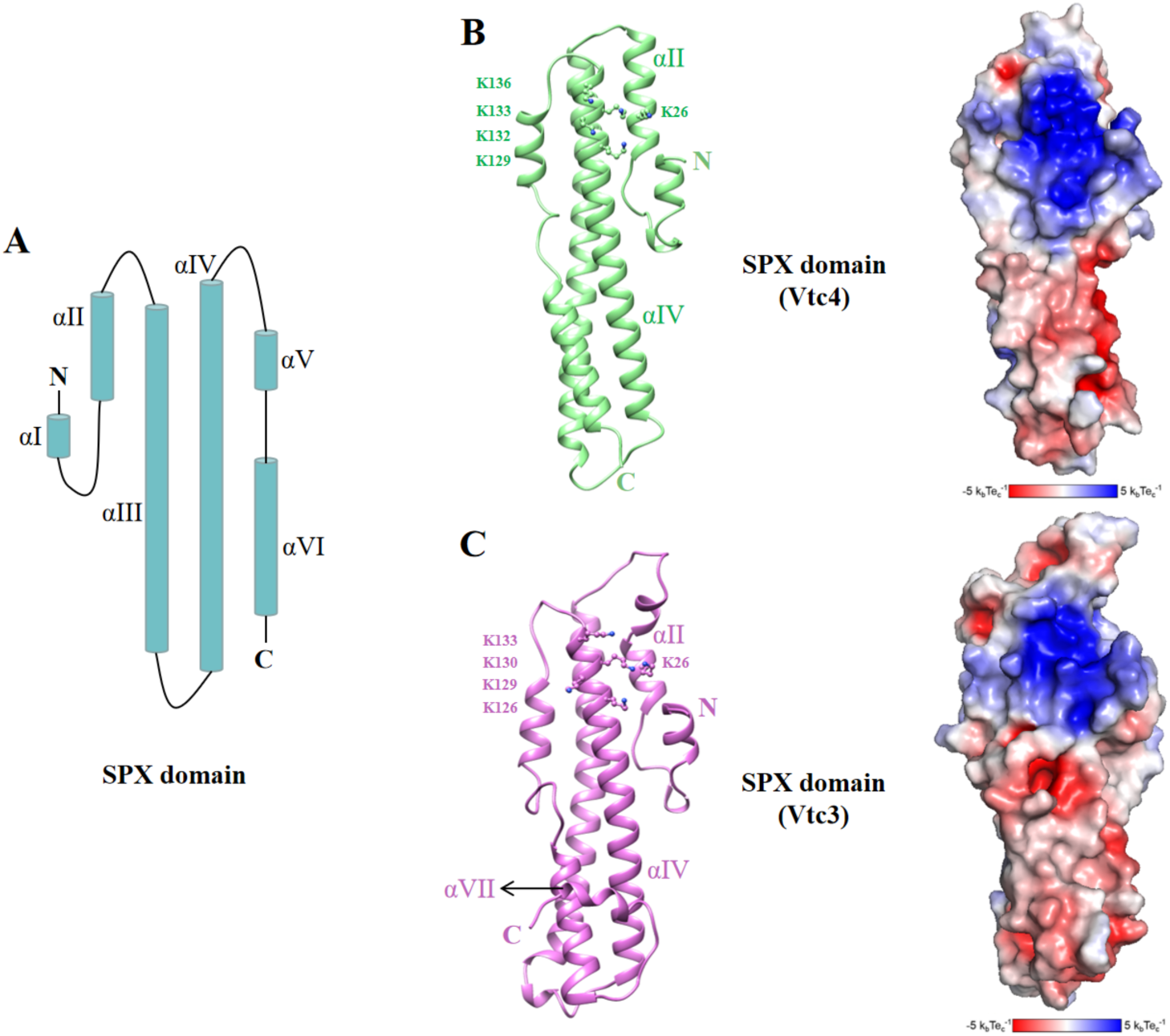
Structure of the SPX domain of Vtc3 and the SPX domain of Vtc4. (A) Schematic representation of the SPX domain. The SPX domain consists of six α- helices, α I-α VI. (B) Structure and electrostatic surface potential of the SPX domain of Vtc4. Key amino acids on the basic surface are highlighted. (C) Structure and electrostatic surface potential of the SPX domain of Vtc3. Key amino acids on the basic surface are highlighted.

**Figure S13.**
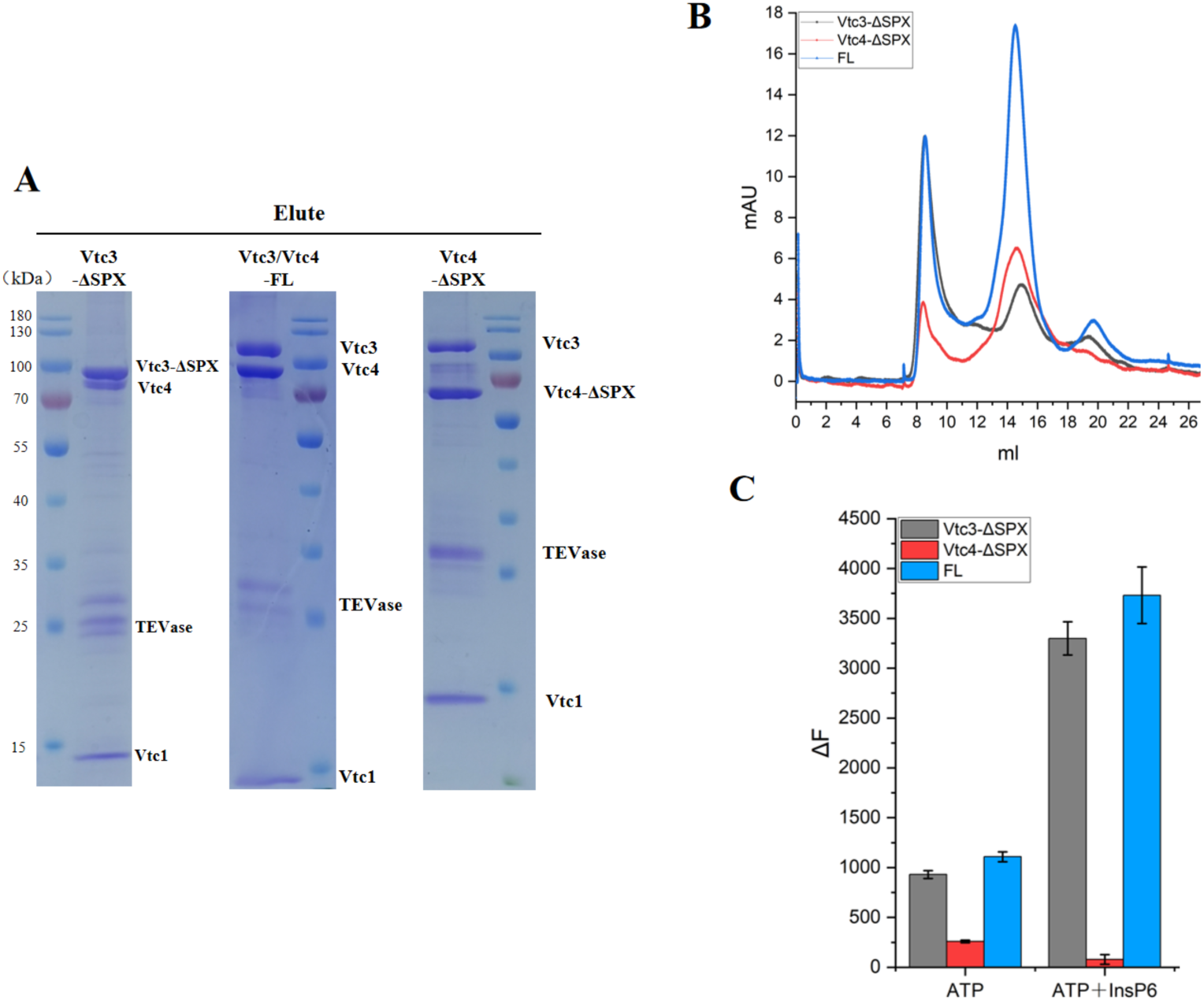
Purification of truncated Vtc4(ΔSPX)/Vtc3/Vtc1 complex and Vtc4/Vtc3(ΔSPX)/Vtc1 complex. (A) The Coomassie blue-stained SDS–PAGE gel of the purified truncated Vtc4(ΔSPX)/Vtc3/Vtc1 complexes, Vtc4/Vtc3(ΔSPX)/Vtc1 complex and Vtc4/Vtc3/Vtc1 complex. (B) Size-exclusion chromatography profile of the truncated Vtc4(ΔSPX)/Vtc3/Vtc1 complex, Vtc4/Vtc3(ΔSPX)/Vtc1 complex and Vtc4/Vtc3/Vtc1 complex. (C) Purified truncated Vtc4(ΔSPX)/Vtc3/Vtc1 complex, Vtc4/Vtc3(ΔSPX)/Vtc1 complex and Vtc4/Vtc3/Vtc1 complex synthesize polyP in the absence or presence InsP6 *in vitro*. The reaction system is detailed in Methods. Data show the mean±s.d (n=3).

**Figure S14.**
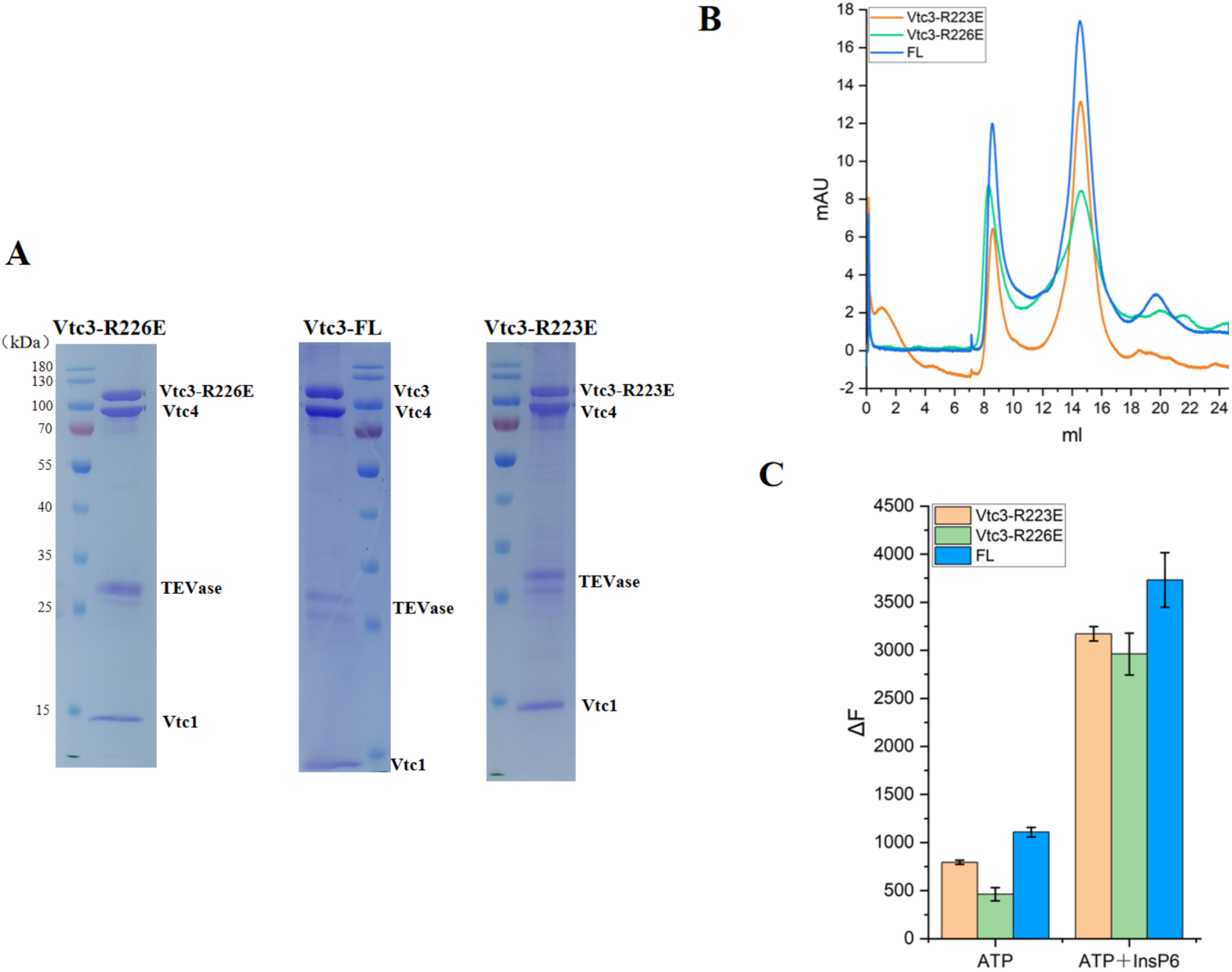
Purification of mutant Vtc4/Vtc3(R223E)/Vtc1 complex and Vtc4/Vtc3(R226E)/Vtc1 complex. (A) The Coomassie blue-stained SDS–PAGE gel of the mutant Vtc4/Vtc3(R223E)/Vtc1 complex, Vtc4/Vtc3(R226E)/Vtc1 complex and Vtc4/Vtc3/Vtc1 complex. (B) Size-exclusion chromatography profile of the mutant Vtc4/Vtc3(R223E)/Vtc1 complex, Vtc4/Vtc3(R226E)/Vtc1 complex and Vtc4/Vtc3/Vtc1 complex. (C) Purified mutant Vtc4/Vtc3(R223E)/Vtc1 complex, Vtc4/Vtc3(R226E)/Vtc1 complex and Vtc4/Vtc3/Vtc1 complex synthesize polyP in the absence or presence InsP6 *in vitro*. The reaction system is detailed in Methods. Data show the mean±s.d (n=3).

**Figure S15.**
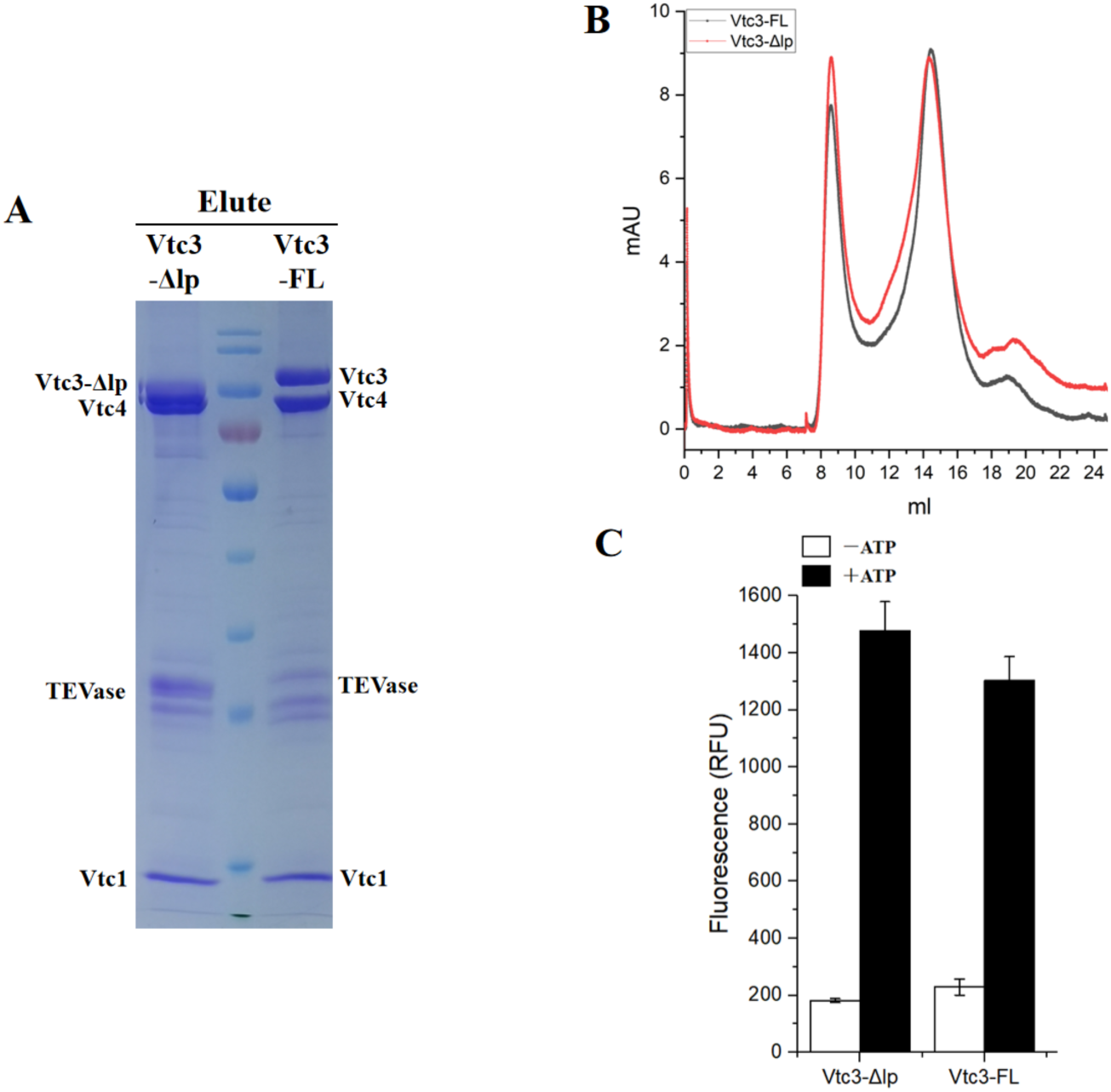
Purification of truncated Vtc4/Vtc3(Δlp)/Vtc1 complexes. (A) The Coomassie blue-stained SDS–PAGE gel of the purified truncated Vtc4/Vtc3(Δlp)/Vtc1 complex and Vtc4/Vtc3/Vtc1 complex. The molecular weights of the marker bands from top to bottom are 180kDa, 130kDa, 100kDa, 70kDa, 55kDa, 40kDa, 35kDa, 25kDa, 15kDa, 10kDa. (B) Size-exclusion chromatography profile of the truncated Vtc4/Vtc3(Δlp)/Vtc1 complex and Vtc4/Vtc3/Vtc1 complex. (C) Purified truncated Vtc4/Vtc3(Δlp)/Vtc1 complexes synthesize polyP in the absence or presence ATP *in vitro*. The reaction system is detailed in Methods. Data show the mean±s.d (n=3).

**Figure S16.**
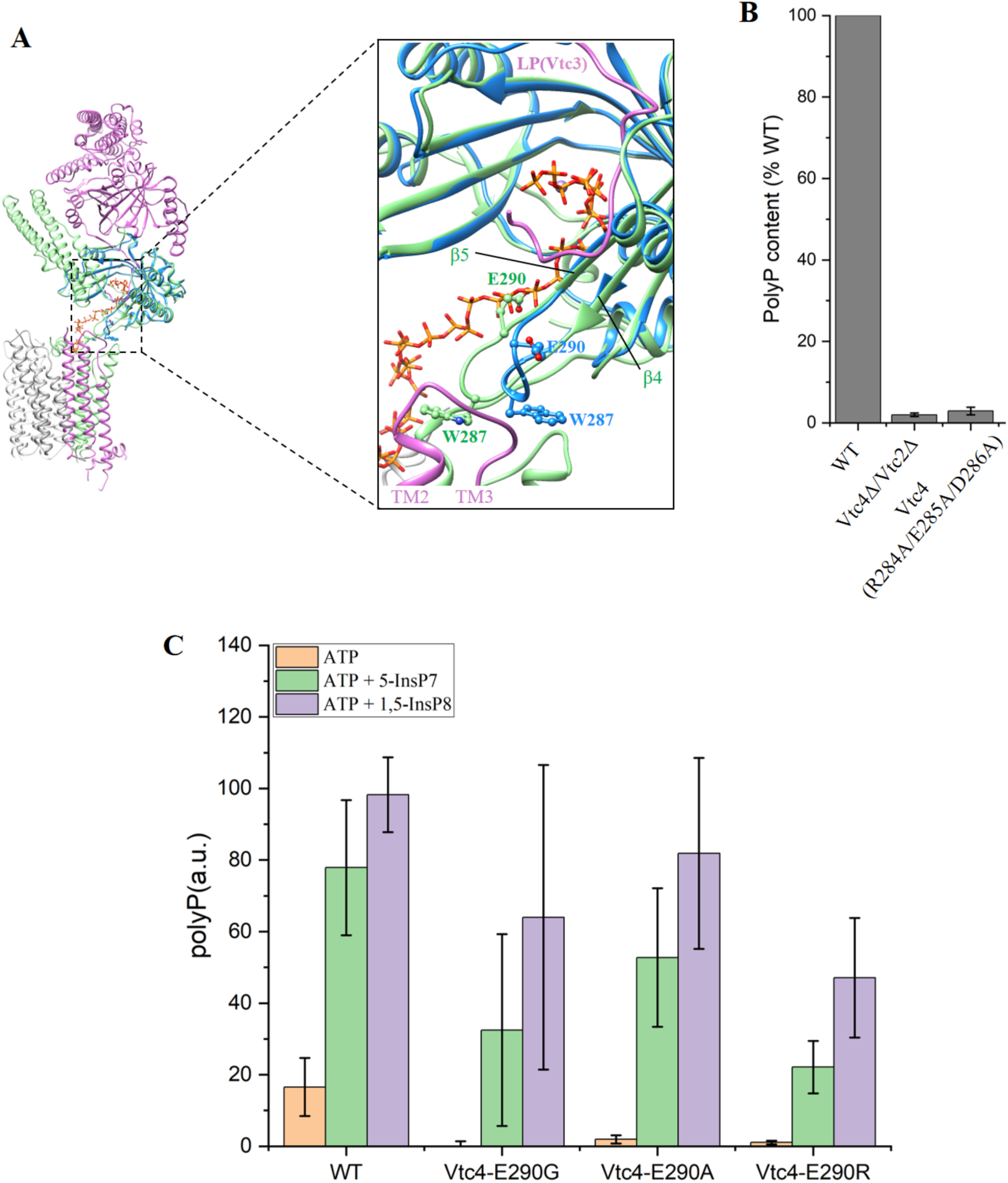
Superposition of the central domain of Vtc4 and the central domain of the polyP-bound Vtc4(PDB: 3G3Q) structures. (A) The structure of the central domain of polyP-bound Vtc4 is shown in blue. The polyP chains are shown in orange to overlap the triphosphates. (B) Cellular polyP content with Vtc4p point mutants expressed under the control of its native promoter in the *vtc4Δ* background. Δ indicates that the entire subunit was knocked out. Data show the mean±s.d (n=3). PolyP synthesis by isolated vacuoles carrying Vtc4(E290G)/Vtc3/Vtc1 complex, Vtc4(E290A)/Vtc3/Vtc1 complex,Vtc4(E290R)/Vtc3/Vtc1 complex, or Vtc4/Vtc3/Vtc1 complex in the absence or presence of 1 μM 5-IP7 or 1,5-IP8 *in vitro*. The reaction system is detailed in Methods. Data show the mean±s.d (n=3).

**Figure S17.**
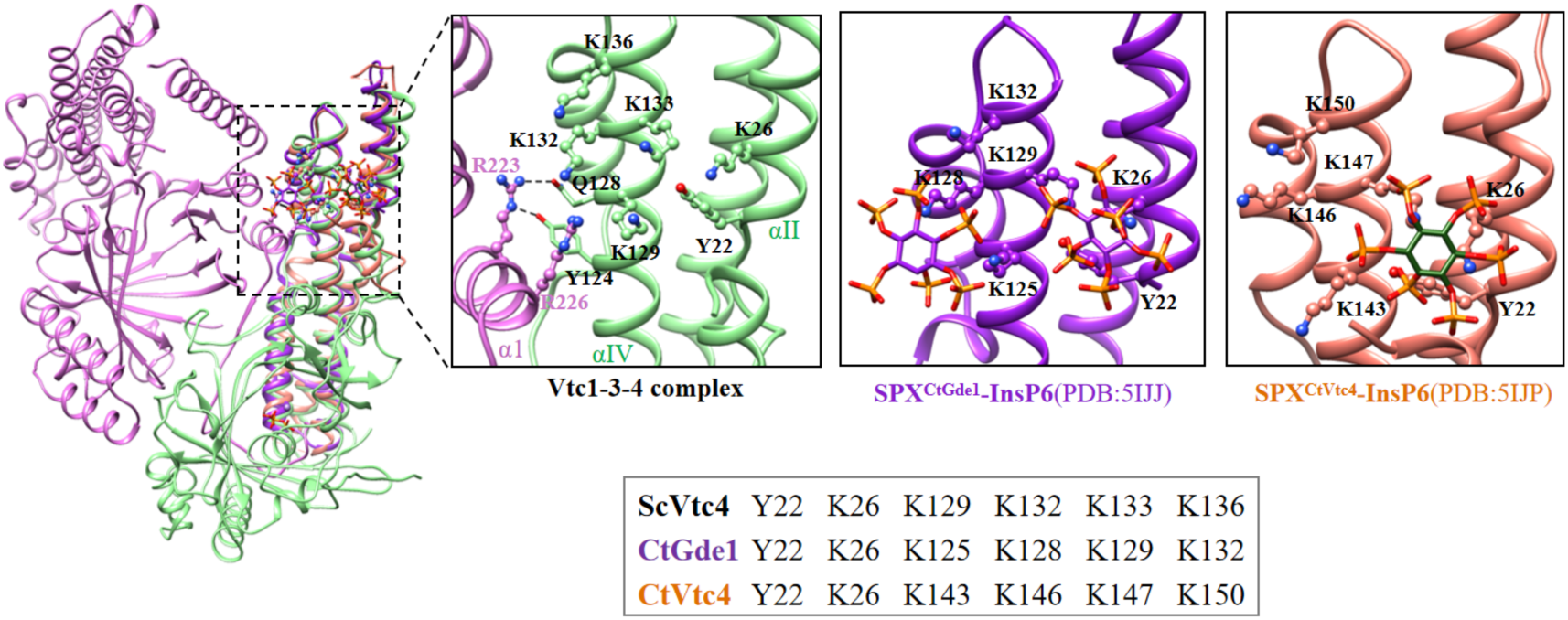
Superimposition of the SPX^CtGde1^-InsP6 (PDB: 5IJJ), the SPX^CtVtc4^- InsP6 (PDB: 5IJP) and the SPX domian of Vtc4. The black squares are the conserved lysines that make up the basic surface.

**Table S1.** Cryo-EM data collection, refinement and validation statistics.

|  | VTC4/3/1 complex<br>(EMDB )<br>(PDB ) |
| --- | --- |
| **Data collection and processing** |  |
| Magnification | 105,000 |
| Voltage (kV) | 300 |
| Electron exposure (e<sup>-</sup>/Å<sup>2</sup>) | 54 |
| Defocus range (μm) | -1.0 ~ -1.5 |
| Pixel size (Å) | 0.851 |
| Symmetry imposed | C1 |
| Initial particle projections (no.) | 3,406,278 |
| Final particle projections (no.) | 1,042,873 |
| Map resolution (Å) | 3.06 |
| FSC threshold | 0.143 |
| Map resolution range (Å) | 2.9-9.8 |
| **Refinement** |  |
| Initial model used | AlphaFold2 |
| Model resolution (Å) | 3.1 |
| FSC threshold | 0.5 |
| Map sharpening B factor (Å<sup>2</sup>) | -97.4 |
| Model composition |  |
| Non-hydrogen atoms | 13,166 |
| Protein residues | 1,594 |
| Ligand | 6 |
| *B*-factors (Å<sup>2</sup>) |  |
| Protein | 0.81/113.97/54.94 |
| Ligand | 34.06/93.82/78.03 |
| R.m.s. deviations |  |
| Bond lengths (Å) | 0.002 |
| Bond angles (°) | 0.493 |
| Validation |  |
| MolProbity score | 1.27 |
| Clashscore | 4.28 |
| Rotamer outliers (%) | 0.00 |
| CaBLAM outliers (%) | 1.68 |
| Ramachandran plot |  |
| Favored (%) | 97.71 |
| Allowed (%) | 2.29 |
| Disallowed (%) | 0.00 |

**Table S2.** Strains used in this study.

| Strain | Relevant genotype | Source |
| --- | --- | --- |
| BJ2168 | MATa: leu2-3, trp1-289, ura3-52, prb1-1122, pep4-3, prc1-407, gal2 | Laboratory |
| BJ2168 *Vtc1Δ* | VTC1::kanMX | This study |
| BJ2168 *Vtc2Δ* | VTC2::kanMX | This study |
| BJ2168 *Vtc3Δ* | VTC3::kanMX | This study |
| BJ2168 *Vtc2Δ Vtc3Δ* | VTC2::URA3 VTC3::kanMX | This study |
| BJ2168 *Vtc4Δ* | VTC4::kanMX | This study |
| BJ2168 *Vtc2*-TAPm | VTC2-TAPm-kanMX | This study |
| BJ2168 *Vtc3*-TAPm | VTC3-TAPm-kanMX | This study |
| BJ2168 *Vtc2*-TAPm *Vtc3*-Strep | VTC2-TAPm-kanMX VTC3-Strep-URA3 | This study |
| BJ2168 *Vtc3*-TAPm *Vtc2*-Strep | VTC3-TAPm-kanMX VTC2-Strep-URA3 | This study |
| BJ2168 *Vtc3*(ΔC24)-TAPm | VTC3(ΔC24)-TAPm-kanMX | This study |
| BJ2168 *Vtc3*(ΔC24)-TAPm *Vtc2Δ* | VTC3(ΔC24)-TAPm-kanMX VTC2::URA3 | This study |
| BJ2168 *Vtc3*(Δlp)-TAPm | VTC3(Δlp)-TAPm-URA3 | This study |
| BJ2168 *Vtc3*(ΔSPX)-TAPm | VTC3(ΔSPX)-TAPm-URA3 | This study |
| BJ2168 *Vtc3*-TAPm *Vtc4*(ΔSPX) | VTC3-TAPm-LEU2 VTC4(ΔSPX)-URA3 | This study |
| BJ2168 *Vtc3*(R223E)-TAPm | VTC3(R223E)-TAPm-URA3 | This study |
| BJ2168 *Vtc3*(R226E)-TAPm | VTC3(R226E)-TAPm-URA3 | This study |
| BJ2168 *Vtc1*(K24E) | VTC1::kanMX VTC1(K24E)-URA3 | This study |
| BJ2168 *Vtc1*(E30A) | VTC1::kanMX VTC1(E30A)-URA3 | This study |
| BJ2168 *Vtc1*(E30R) | VTC1::kanMX VTC1(E30R)-URA3 | This study |
| BJ2168 *Vtc1*(R31E) | VTC1::kanMX VTC1(R31E)-URA3 | This study |
| BJ2168 *Vtc1*(R83A) | VTC1::kanMX VTC1(R83A)-URA3 | This study |
| BJ2168 *Vtc1*(R83E) | VTC1::kanMX VTC1(R83E)-URA3 | This study |
| BJ2168 *Vtc3*(K698E) *Vtc2Δ* | VTC3::kanMX VTC3(K698E)-LEU2 VTC2::URA3 | This study |
| BJ2168 *Vtc3*(V699D) *Vtc2Δ* | VTC3::kanMX VTC3(V699D)-LEU2 VTC2::URA3 | This study |
| BJ2168 *Vtc3*(E704A) *Vtc2Δ* | VTC3::kanMX VTC3(E704A)-LEU2 VTC2::URA3 | This study |
| BJ2168 *Vtc3*(E704R) *Vtc2Δ* | VTC3::kanMX VTC3(E704R)-LEU2 VTC2::URA3 | This study |
| BJ2168 *Vtc1*(R705E) *Vtc2Δ* | VTC3::kanMX VTC3(R705E)-LEU2 VTC2::URA3 | This study |
| BJ2168 *Vtc3*(R762A) *Vtc2Δ* | VTC3::kanMX VTC3(R762A)-LEU2 VTC2::URA3 | This study |
| BJ2168 *Vtc3*(R762E) *Vtc2Δ* | VTC3::kanMX VTC3(R762E)-LEU2 VTC2::URA3 | This study |
| BJ2168 *Vtc3*(L765D) *Vtc2Δ* | VTC3::kanMX VTC3(L765D)-LEU2 VTC2::URA3 | This study |
| BJ2168 *Vtc3*(L774D) *Vtc2Δ* | VTC3::kanMX VTC3(L774D)-LEU2 VTC2::URA3 | This study |
| BJ2168 *Vtc4*(ΔHH) | VTC4::kanMX VTC4(ΔHH)-URA3 | This study |
| BJ2168 *Vtc4*(R196E) | VTC4::kanMX VTC4(R196E)-URA3 | This study |
| BJ2168 *Vtc4*(K200A) | VTC4::kanMX VTC4(K200A)-URA3 | This study |
| BJ2168 *Vtc4*(R253E) | VTC4::kanMX VTC4(R253E)-URA3 | This study |
| BJ2168 *Vtc4*(K256E) | VTC4::kanMX VTC4(R256E)-URA3 | This study |
| BJ2168 *Vtc4*(R264A) | VTC4::kanMX VTC4(R264A)-URA3 | This study |
| BJ2168 *Vtc4*(R266A) | VTC4::kanMX VTC4(R266A)-URA3 | This study |
| BJ2168 *Vtc4*(R264A/R266A) | VTC4::kanMX VTC4(R264A/R266A)-URA3 | This study |
| BJ2168 *Vtc4*(K281A) | VTC4::kanMX VTC4(K281A)-URA3 | This study |
| BJ2168 Vtc4(W287D) | VTC4::kanMX VTC4(W287D)-URA3 | This study |
| BJ2168 Vtc4(K291E) | VTC4::kanMX VTC4(K291E)-URA3 | This study |
| BJ2168 Vtc4(K294E) | VTC4::kanMX VTC4(K294E)-URA3 | This study |
| BJ2168 Vtc4(Y359F) | VTC4::kanMX VTC4(Y359F)-URA3 | This study |
| BJ2168 Vtc4(R361A) | VTC4::kanMX VTC4(R361A)-URA3 | This study |
| BJ2168 Vtc4(R373E) | VTC4::kanMX VTC4(R373E)-URA3 | This study |
| BJ2168 Vtc4(E426A) | VTC4::kanMX VTC4(E426A)-URA3 | This study |
| BJ2168 Vtc4(K428E) | VTC4::kanMX VTC4(K428E)-URA3 | This study |
| BJ2168 Vtc4(K455E) | VTC4::kanMX VTC4(K455E)-URA3 | This study |
| BJ2168 Vtc4(S457A) | VTC4::kanMX VTC4(S457A)-URA3 | This study |
| BJ2168 Vtc4(K458L) | VTC4::kanMX VTC4(K458L)-URA3 | This study |
| BJ2168 Vtc4(K458I) | VTC4::kanMX VTC4(K458I)-URA3 | This study |
| BJ2168 Vtc4(K458A) | VTC4::kanMX VTC4(K458A)-URA3 | This study |
| BJ2168 Vtc4(R618E) | VTC4::kanMX VTC4(R618E)-URA3 | This study |
| BJ2168 Vtc4(P621D) | VTC4::kanMX VTC4(P621D)-URA3 | This study |
| BJ2168 Vtc4(K622E) | VTC4::kanMX VTC4(K622E)-URA3 | This study |
| BJ2168 Vtc4(E628A) | VTC4::kanMX VTC4(E628A)-URA3 | This study |
| BJ2168 Vtc4(E628R) | VTC4::kanMX VTC4(E628R)-URA3 | This study |
| BJ2168 Vtc4(R629E) | VTC4::kanMX VTC4(R629E)-URA3 | This study |
| BJ2168 Vtc4(R681A) | VTC4::kanMX VTC4(R681A)-URA3 | This study |
| BJ2168 Vtc4(R681E) | VTC4::kanMX VTC4(R681E)-URA3 | This study |

TAPm is a modified TAP tag consisting of 6His-TEV-Protein A.

